# Nucleoside Diphosphate Kinases 1 and 2 regulate a protective liver response to a high-fat diet

**DOI:** 10.1101/2023.01.15.524116

**Authors:** Domenico Iuso, Isabel Garcia-Saez, Yohann Couté, Yoshiki Yamaryo-Botté, Elisabetta Boeri Erba, Annie Adrait, Nour Zeaiter, Malgorzata Tokarska-Schlattner, Zuzana Macek Jilkova, Fayçal Boussouar, Sophie Barral, Luca Signor, Karine Couturier, Azadeh Hajmirza, Florent Chuffart, Anne-Laure Vitte, Lisa Bargier, Denis Puthier, Thomas Decaens, Sophie Rousseaux, Cyrille Botté, Uwe Schlattner, Carlo Petosa, Saadi Khochbin

**Affiliations:** CNRS UMR 5309/INSERM U1209/Université Grenoble-Alpes/Institute for Advanced Biosciences, 38706, La Tronche, France; Univ. Grenoble Alpes, CNRS, CEA, Institut de Biologie Structurale (IBS), 38000 Grenoble, France; Université Grenoble Alpes, INSERM, CEA, UMR BioSanté U1292, CNRS, CEA, FR2048, 38000 Grenoble, France; Université Grenoble-Alpes, INSERM, Laboratory of Fundamental and Applied Bioenergetics, Grenoble, France.; CHU Grenoble Alpes, Service d’hépato-gastroentérologie, Pôle Digidune, 38700 La Tronche, France; Aix Marseille Université, INSERM, TAGC, TGML, 13288, Marseille, France; Université Grenoble-Alpes, INSERM, Institut Universitaire de France, Laboratory of Fundamental and Applied Bioenergetics, Grenoble, France.

## Abstract

*De novo* lipogenesis (DNL), the process whereby cells synthesize fatty acids from acetyl-coenzyme A (acetyl-CoA), is deregulated in diverse pathologies, including cancer. Here we report that DNL is negatively regulated by Nucleoside Diphosphate Kinases 1 and 2 (NME1/2), housekeeping enzymes involved in nucleotide homeostasis that were recently discovered to bind co-enzyme A (CoA). We show that NME1 additionally binds acetyl-CoA and that ligand recognition involves a unique binding mode dependent on the CoA/acetyl-CoA 3’ phosphate. We report that *Nme2* knockout mice fed a high-fat diet (HFD) exhibit excessive triglyceride synthesis and liver steatosis. In liver cells NME2 mediates a gene transcriptional response to HFD leading to DNL repression and activation of a protective gene expression program via targeted histone acetylation. Our findings implicate NME1/2 in the epigenetic regulation of a protective liver response to HFD and suggest a potential role in controlling acetyl-CoA usage between the competing paths of histone acetylation and DNL.

## INTRODUCTION

De novo lipogenesis (DNL) is the pathway used primarily by the liver and adipose tissue to synthesize fatty acids from excess carbohydrates or other precursors^1^. Deregulated DNL, especially in the liver, is implicated in diverse pathologies^2^. Increased rates of DNL are associated with nonalcoholic fatty liver disease^3, 4^, insulin resistance and type 2 diabetes^4, 5^, cardiovascular disease^6^, incident heart failure^7^ and cancer^8^. The main carbon source for DNL is the cytoplasmic pool of acetyl coenzyme A (AcCoA), which is derived from either citrate or acetate through the activities of ATP Citrate Lyase (ACLY) and AcCoA synthetase (ACSS2), respectively^9, 10^ (**Fig. 1a**). AcCoA is irreversibly converted to malonyl-CoA by the action of acetyl-CoA carboxylases 1 and 2 (ACC1 and ACC2), which localize to the cytosol and outer mitochondrial membrane, respectively. While the malonyl-CoA produced by ACC2 negatively regulates fatty acid import into mitochondria and hence β-oxidation, that produced by ACC1 is used in iterative cycles of fatty acid elongation by fatty acid synthase (FAS) to generate fatty acids, predominantly the C16:0 fatty acid palmitate.

**Figure 1.**
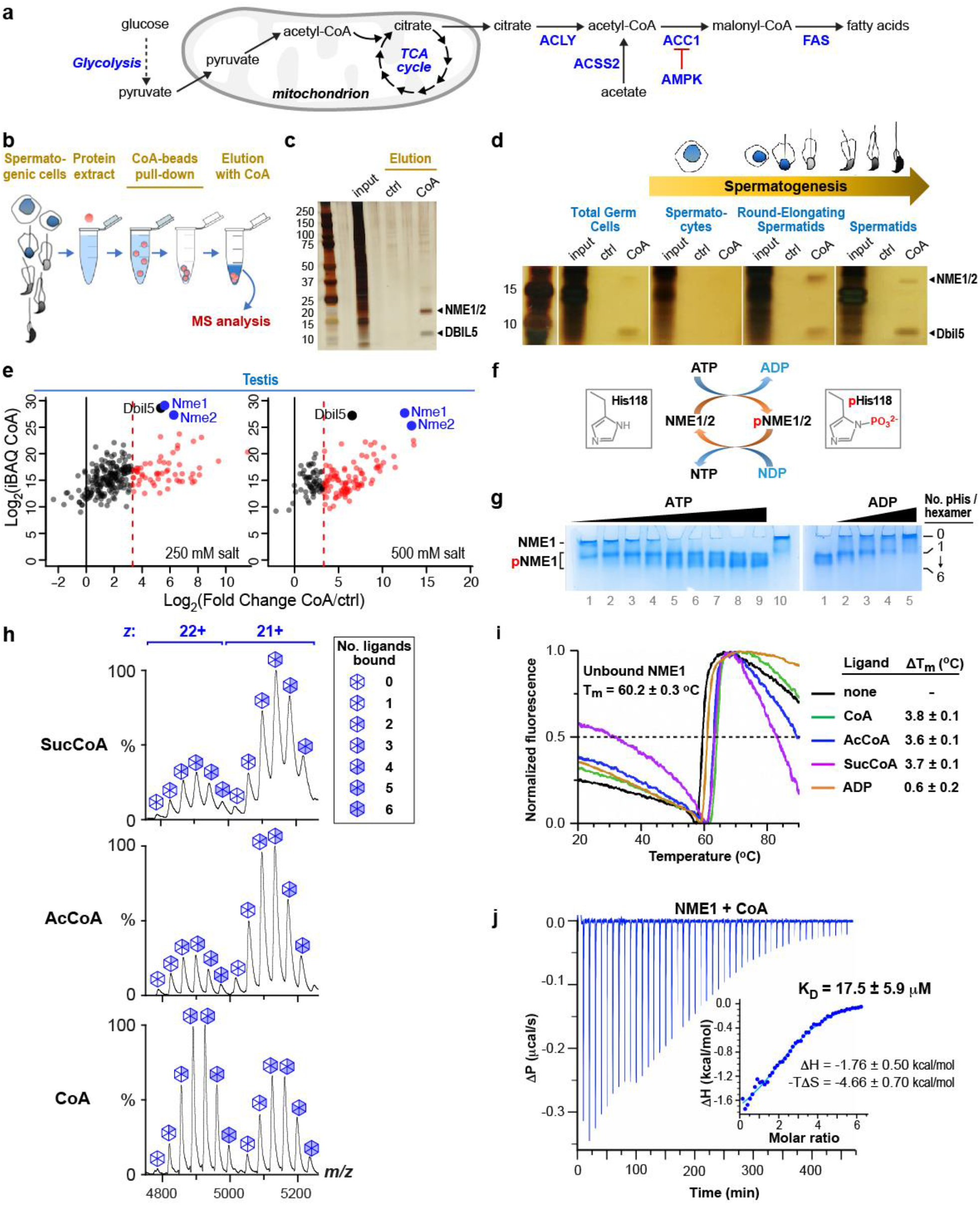
NME1/2 are major CoA-binding factors in mouse spermatogenic cells and bind short- chain acyl-CoA ligands *in vitro*. **a.** Overview of DNL. Under conditions of carbohydrate excess, mitochondrial citrate produced by the tricarboxylic acid (TCA) cycle is exported to the cytosol and converted to AcCoA by ACLY. Acetate synthesized from pyruvate by both enzymatic and non-enzymatic means^66^ is converted to AcCoA by ACSS2. The commitment step of DNL catalysed by ACC1 is negatively regulated by AMPK. **b.** Strategy for purifying CoA-binding factors present in extracts from total testis or fractionated spermatogenic cells. After the incubation of extracts with CoA-coated Sepharose beads and successive washes, bound proteins were eluted with either free CoA or water as a control. **c.** Silver-stained electrophoresis gel of proteins eluted from CoA beads with either water (ctrl) or CoA. The major proteins captured by CoA are indicated. **d.** Total testis extracts or equivalent amounts of extracts from the indicated fractionated spermatogenic cells were incubated with CoA beads and processed as in (**b**). **e.** The identities and abundances of proteins eluted by free CoA (CoA) and water (Ctrl) were deduced from MS-based proteomic analyses. The graphs display the log_2_(iBAQ) of the proteins in CoA (representative of the relative abundance of the different proteins identified in this sample) on the *y*-axis plotted against the log_2_(Fold Change CoA/Ctrl) on the *x*-axis (representative of the differential abundance between CoA and Ctrl fishing). For each representation, the black and dashed red vertical lines indicate respectively log_2_(Fold Change) = 0, and log2(Fold Change) = 3.32 (i.e. Fold Change (CoA/Ctrl) = 10). Black and red dots highlight proteins with a Fold Change (CoA/Ctrl) < 10 and ≥ 10, respectively. The three most abundant proteins, NME1/2 and DBIL5, are indicated. CoA pull-downs were performed on total testis extracts in the presence of either 250 (left) or 500 (right) mM KCL. **f.** Scheme illustrating phosphohistidine-mediated NDPK activity. Only one direction of the fully reversible reaction is shown. **g.** Gel shift experiments confirm NDPK activity of purified recombinant NME1. *Left gel:* NME1 is phosphorylated by ATP. Coomassie-stained native polyacrylamide gel of bacterially expressed NME1 incubated without (lane 10) or with (lanes 1-9) increasing amounts of ATP. At higher ATP concentrations more subunits within each NME1 hexamer are phosphorylated, resulting in additional higher mobility bands on the gel. *Right gel:* NME1 is dephosphorylated by ADP. Native gel of phosphorylated NME1 incubated without (lane 1) or with (lanes 2-5) increasing amounts of ADP, illustrating phosphoryl group transfer from NME1 to ADP. **h.** Native MS analysis of recombinant purified NME1 incubated with CoA, AcCoA or SucCoA, as indicated. Peaks corresponding to charge states +22 and +21 represent the NME1 hexamer with 0 to 6 ligands bound. **i.** Thermal denaturation profile of NME1 measured by nanoDSF. Representative profiles are shown for NME1 in the absence and presence of CoA ligands. *T*_m_ and Δ*T*_m_ values represent the mean and SD from three independent experiments. **j.** Representative ITC profile of CoA binding by NME1. Differential power (Δ*P*) data time course of raw injection heats for a titration of CoA into NME1. *Inset:* Normalized binding enthalpies corrected for heat of dilution as a function of binding site saturation. Data were fit using a single- site binding model. *K*_D_ and thermodynamic parameters represent mean and SD values determined from three independent experiments.

ACC1 catalyses the first committed step of DNL and hence is subject to tight regulation. Besides allosteric activation by citrate and product inhibition by malonyl- and palmitoyl-CoA, ACC1 is also tightly regulated by the cellular energy level. Energy deficiency, characterized by a depletion of ATP and an accumulation of ADP and AMP, triggers the inactivating phosphorylation of ACC1 by AMP-activated protein kinase (AMPK), thereby inhibiting DNL. High levels of ATP, in contrast, favour the massive use of AcCoA in fatty acid synthesis. This well-characterized mechanism establishes a tight relationship between ATP levels, ACC1 activity and fatty acid synthesis and storage^11^. Here we report the discovery of an additional layer of DNL regulation, in which the key players are the nucleoside diphosphate kinases (NDPKs) 1 and 2 (NME1 and NME2). NDPKs mediate a large variety of cellular functions that include nucleotide homeostasis, endocytosis, intracellular trafficking, cell motility and DNA repair, with implications in cancer and metastatic cancer development^12^. In mammals the most abundant and best studied NDPKs are NME1 and NME2 (hereafter abbreviated NME1/2), which are closely related isoforms (88% sequence identity) with similar functional attributes and localization in the cytosol and nucleus. The best-characterized activity of these enzymes is the phosphorylation of nucleoside diphosphates (NDPs), using nucleoside triphosphates (NTPs), primarily ATP, as a donor. This reaction, which plays a key role in equilibrating the nucleotide pools in the cell, proceeds by a ping-pong mechanism involving the formation of a phosphohistidine intermediate in the catalytic site. NME1/2 can also transfer the phosphate group from this intermediate to a histidine in unrelated substrate proteins and accordingly are also described as histidine kinases^13^.

Recently, NME1/2 were reported to have the remarkable and unexpected ability to bind CoA^14^. CoA binds non-covalently to NME1 and competitively inhibits NDPK activity. Moreover, in cells under oxidative or metabolic stress, NME1 undergoes CoAlation (covalent modification via disulfide formation with the CoA thiol group) at a specific cysteine near the enzyme’s active site, resulting in the stable inhibition of NDPK activity^14^. A more recent study reported that NME1/2 also interact with long-chain fatty acyl (LCFA)-CoA, which binds via its nucleotide moiety to the enzyme’s active site, as revealed by the crystal structure of NME2 bound to myristoyl-CoA^15^. The study also showed that the increased production of cellular LCFA-CoA inhibited clathrin-mediated endocytosis, an NME1/2-dependent process, and that LCFA-CoA compromised the metastasis suppressor function of NME1 in mouse models of breast cancer under high-fat diet (HFD) conditions^15^. Taken together, these studies suggest that CoA and its derivatives may regulate diverse NME1/2-mediated processes.

Here we report a previously undescribed role for NME1/2 in the control of DNL as well as in a protective liver cell response to a HFD challenge. We present *in vitro* and structural data showing that, in addition to CoA and LCFA-CoA, NME1 also binds AcCoA, the most abundant CoA derivative, and that ligand recognition occurs via a unique nucleotide binding mode critically dependent on the CoA 3’ phosphate group. Moreover, binding is inhibited by ATP-induced histidine phosphorylation and hence potentially sensitive to the cellular energy status. Focusing on a mouse knockout model with drastically reduced NME1/2 levels, we show that the major cytoplasmic AcCoA consuming pathway is stimulated during liver fatty acid synthesis, leading to increased liver triglyceride levels and liver steatosis. Under HFD challenge, AcCoA binding by NME1/2 exerts an inhibitory role on ACC1-dependent fatty acid synthesis, specifically repressing genes encoding key transcription factors involved in lipogenesis and fatty acids metabolism. At the same time, NME1/2 mediate an increased targeted histone H3K9 acetylation, activating a gene signature known to protect liver cells during regeneration. These observations identify NME1/2 as a critical regulator of the competing processes of histone acetylation and DNL, placing NME1/2 among a select group of metabolic enzymes with a moonlighting function in epigenetic regulation^16^.

## RESULTS

### NME1/2 are major CoA-binding factors in mouse spermatogenic cells

NME1/2 were previously identified as CoA or LCFA-CoA interacting proteins in HEK293 cell lines and rat heart extracts^14, 15^. We independently discovered that NME1/2 bind CoA while investigating the global genome reprogramming that occurs in mouse spermatogenic cells during post-meiotic (spermatid) maturation, which is characterized by a switch from a nucleosome- to a protamine-based organization following the nearly genome-wide eviction of histones. Surmising that the genome-wide increase in histone acetylation observed prior to histone eviction^17–19^ requires an enhanced production, management and use of AcCoA, we sought to identify CoA- binding regulatory factors involved in managing the AcCoA pool in spermatids. We incubated CoA-coated beads with extracts from mouse testis and spermatogenic cells enriched at different stages of their development by fractionation and subsequently eluted CoA-binding proteins with free CoA (**Fig. 1b**). Mass spectrometry (MS) based proteomic analysis identified several CoA-associated proteins common to all samples, including total testis extracts pulled down with different stringencies as well as extracts from fractionated spermatogenic cells (**Table S1**). In addition to the known testis-specific acyl-CoA binding protein DBIL5^20^, the most prominent CoA-binding factors were NME1/2 (**Fig. 1c-e**, **Fig. S1** and **Table S1**). This is consistent with the strong prevalence of NME1/2 detected among CoA-binding factors in diverse somatic rat tissues^14^, highlighting the ubiquitous nature of NME1/2’s CoA-binding functionality.

### NME1 binds acetyl-CoA *in vitro*

Since NME1/2 recognizes myristoyl-CoA exclusively by its nucleotide moiety^15^, NME1/2 should also bind AcCoA and other short-chain acyl-CoA molecules such as succinyl-CoA (SucCoA). To verify this we assessed the ability of purified NME1 to bind these molecules *in vitro*. NME1/2 are hexameric enzymes that catalyze the reaction XTP + YDP Δ XDP + YTP by first transferring the γ- phosphoryl group from XTP (usually ATP) to residue His118, and then from the phosphohistidine to YDP, where Y is any of the four common (deoxy)nucleosides (**Fig. 1f**). We confirmed that recombinant murine NME1 purified from bacteria could be phosphorylated by ATP and subsequently dephosphorylated by ADP or GDP (**Fig. 1g** and **Fig. S2**). NME1 incubated with CoA was then analysed by native MS, which allows the mass measurement of intact non-covalent complexes^21^. NME1 hexamers were observed in the unbound state and bound to a variable number of CoA molecules ranging from 1 to 6 (**Fig. 1h**). The observed pattern of peak intensities matched the distribution of ligand-bound states expected for a hexamer with six independent binding sites that are each approximately half occupied (**Fig. S3**). Incubating NME1 with either AcCoA or SucCoA yielded similar spectra, revealing that NME1 does not strongly discriminate between these ligands and CoA.

To further evaluate the relative affinity of NME1 for CoA, AcCoA and SucCoA we used nano-differential scanning fluorimetry (nanoDSF) to assess ligand-induced thermal stabilization. The recorded profiles revealed that the melting temperature of NME1 (*T*_m_=60°C) was enhanced similarly by each of the tested ligands (Δ*T*_m_≍3.7 °C), confirming that AcCoA and SucCoA bind NME1 with similar affinity as CoA (**Fig. 1i**). Since the latter affinity has not previously been quantified we analysed the NME1:CoA interaction by isothermal titration calorimetry (ITC). This revealed a dissociation constant (*K*_d_) of ∼18 μM (**Fig. 1j**), comparable to or lower than the *K*_d_ values of 25-120 μM reported for the binding of NDPKs to ADP^22–25^. Consistent with this observation, ADP was less effective than the CoA ligands at stabilizing NME1 in the nanoDSF assay (Δ*T*_m_=0.6°C, **Fig. 1i**). Taken together, these data indicate that CoA, AcCoA and SucCoA have similar affinity for NME1 and bind at least as well, if not better, than canonical NDP substrates.

### NME1 recognizes CoA via a unique nucleotide binding mode

Efforts to cocrystallize murine NME1 with the above ligands led to high-resolution (1.96 and 2.2 Å, respectively) crystal structures of NME1 bound to SucCoA and ADP and a somewhat lower (2.6 Å) resolution structure bound to CoA (**Table S2**). As expected, the structure with ADP closely matches that of the ADP-bound human ortholog^26^ while those with CoA and SucCoA are nearly identical to each other and closely resemble that of human NME2 bound to myristoyl-CoA^15^ (**Table S3** and **Fig. S4**). Briefly, each NME1 subunit within the D_3_-symmetric hexamer comprises an antiparallel 4-stranded β-sheet flanked on either side by a layer of helices^27^. ADP is sandwiched between the α_A_-α_2_ helical hairpin and an α3-β4 loop segment called the “Kpn loop” that define the enzyme’s active site (**Fig. 2a**). CoA and SucCoA bind to this site via their common nucleotide moiety, the only part of these ligands specifically recognized by NME1. (Hence, the CoA and SucCoA ligands in our structures are hereafter collectively called “CoA” unless otherwise specified). In the electron density maps, no or only poorly defined density is observed for the pantetheine and succinyl moieties of these ligands (**Fig. S5**), except in a few NME1 subunits where they make non-specific contacts with nearby residues. Consequently, the acyl-bearing end of CoA remains solvent accessible when bound to NME1 and potentially free to interact with other protein partners, consistent with our findings that NME1 binds similarly to acylated and non-acylated CoA (**Fig. 1h,i**).

**Figure 2.**
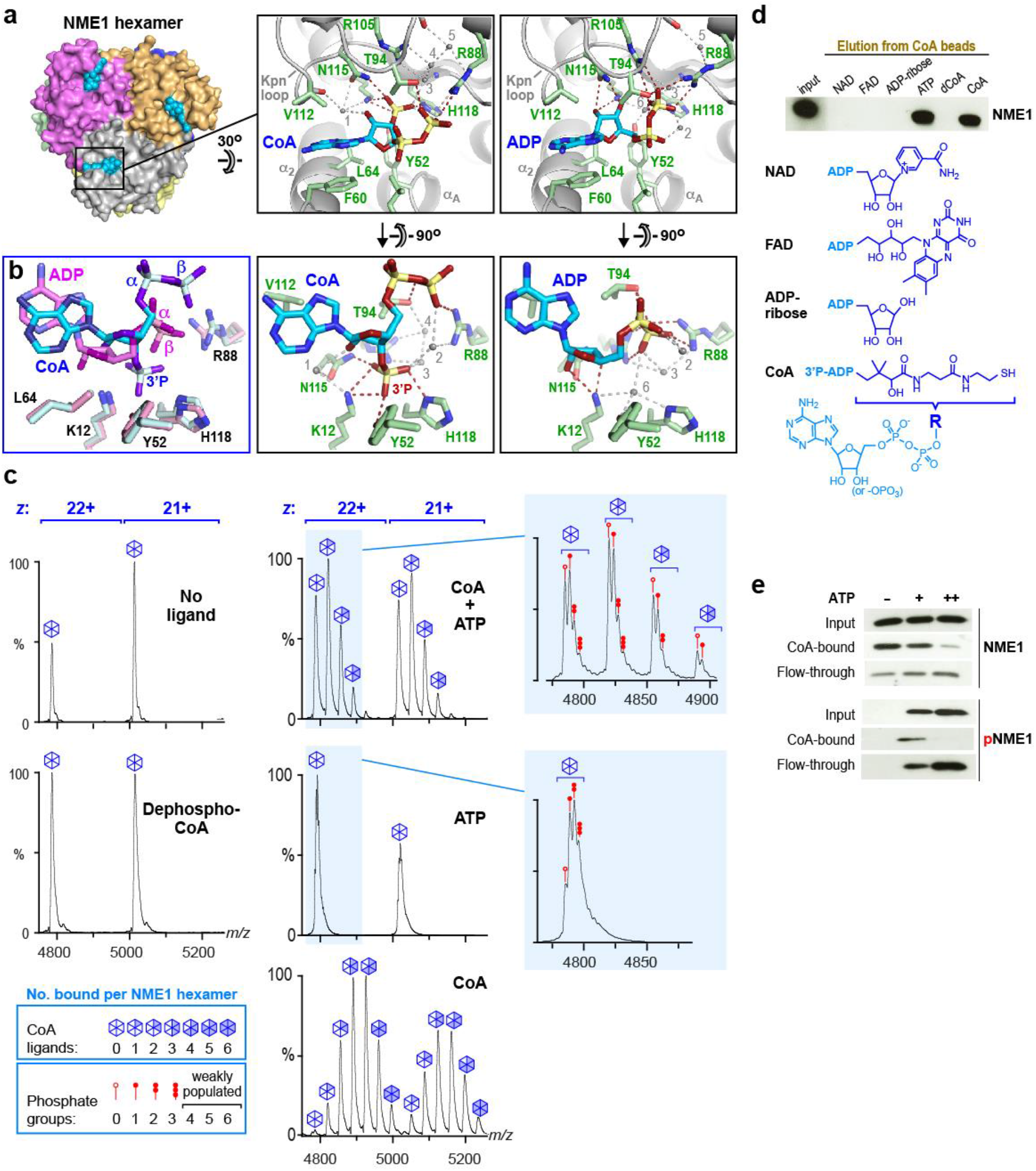
NME1 recognizes CoA via a unique binding mode dependent on the 3’ phosphate and mutually exclusive with histidine phosphorylation. **a.** Structural comparison of NME1 bound to CoA/SucCoA (left) and ADP (right). Hydrogen bonds between protein and ligand atoms are shown as red dashed lines. Water-mediated bonds are shown in gray. Water molecules are indicated as gray spheres. **b.** Superimposition of the CoA and ADP ligands showing the 40° relative rotation of the adenine base and the repositioning of the α- and β-phosphate groups, which are relatively shifted by 2.9 and 3.9 Å, respectively. **c.** Native MS analysis of NME1 incubated in the absence of ligand (upper left) or presence of dephospho-CoA (lower left), CoA (lower right; same spectrum as in Fig. 1h), ATP (middle right) or an equimolar mixture of CoA and ATP (each in 5-fold excess relative to NME1; upper right), which leads to a mixture of CoA binding and histidine phosphorylation. Spectra show that the 3’ phosphate of CoA is critical for binding and that CoA binding and phosphorylation of the NME1 hexamer are mutually competitive. **d.** NME1 does not promiscuouly bind common ADP-containing metabolites. NME1-loaded CoA beads (input) were incubated with an excess of the indicated molecules and the release of NME1 was visualized by an immunoblot of the flow-through probed with an NME1 antibody. dCoA is dephospho-CoA. **e.** NME1 (3 μg) loaded on CoA beads were incubated in the absence or presence of either 30 or 300 μM ATP and the amounts of bound (CoA-bound) and released (flow through) NME1 were visualized by immunoblotting (upper panels). The lower panels show the same blots probed with an anti-phosphohistidine antibody.

How NME1/2 recognizes the CoA nucleotide in atomic detail has not previously been described and hence is presented here. Surprisingly, NME1 recognizes ADP and CoA via completely distinct binding modes (**Fig 2a,b** and **Fig. S6**). Of the 10 direct hydrogen bonds mediating the NME1:ADP interaction, only two are conserved in the NME1:CoA complex. This is striking because the ligand-interacting residues of NME1 adopt nearly identical conformations in both complexes and because, apart from its 3’ phosphate group, the CoA nucleotide is otherwise chemically identical to ADP. The CoA 3’-phosphate group sits next to the catalytic His118 side chain at the bottom of the ligand binding pocket (**Fig. 2a,b**), the same site putatively occupied by the ATP γ phosphate group (**Fig. S7**). As a result, the ribose ring sits 2 Å higher than in the ADP-bound structure, adopting a C_2’_ instead of a C_3’_ *endo* pucker. Compared to ADP, the α- and β- phosphates of CoA are displaced upwards and are solvent accessible, pointing out of the pocket to direct the CoA pantetheine end away from the protein surface. This contrasts with the ADP-bound structure, where the β-phosphate is buried deep inside the protein and folds back towards the ribose 3’ OH group, with which it forms an intramolecular H bond critical for catalysis^28, 29^. Hence, whereas the β-phosphate of ADP is intimately recognized by NME1 via numerous H bonds, that of CoA makes fewer protein contacts (**Fig. S6**) and exhibits high thermal mobility (**Fig. S8a**). The α-phosphate of CoA is also highly mobile, only contacting NME1 through a salt bridge with residue Arg58 in a few hexamer subunits (**Fig. S8b**).

How does NME1 accommodate two very different nucleotide binding modes? First, as in the ADP complex, the adenine base of CoA is sandwiched between hydrophobic residues from the Kpn loop and α_A_-α_2_ helical hairpin but is rotated by ∼40° in the plane of the base to compensate for the shifted ribose ring and maintain a stacking interaction with residue Phe60 (**Fig. 2a,b**). This rotation is possible because NME1 binds nucleotides without forming H-bonds with the base moiety, allowing NME1 to bind substrates regardless of the identity of the base. Second, the high resolution of our NME1/SucCoA crystal structure revealed several water molecules structurally conserved across hexamer subunits that play an important role in compensating for the shift of CoA nucleotide atoms relative to their counterparts in ADP (**Fig. S6**). Finally, the outward shift of the β-phosphate is accompanied by an inward shift of residue Thr94 in the α3-β4 loop that allows it to hydrogen bond with the repositioned β-phosphate (**Fig. 2a** and **Fig. S9a**).

To confirm the non-canonical binding mode of CoA we exploited the shift in Thr94 position to design a mutation that would selectively disrupt CoA binding without completely abolishing the interaction with ADP and ATP. *In silico* modelling suggested that replacing Thr94 by an Asp residue should yield an NME1 mutant that could accommodate ADP, but not CoA, in the active site (**Fig. S9b**), thereby disrupting CoA binding while allowing NDP phosphorylation to continue. Accordingly, we generated the T94D mutant and evaluated it for NDPK and CoA-binding activity. WT and mutant forms of GST-tagged NME1 could both be phosphorylated by ATP and subsequently dephosphorylated by GDP, confirming that the T94D mutant retained significant NDPK activity, albeit at a reduced level compared to the WT (**Fig. S9c** upper panel**, d**). In contrast, whereas WT NME1 bound strongly to immobilized CoA, no interaction was detected between the T94D mutant and the CoA beads, revealing a dramatic loss of CoA-binding capacity (**Fig. S9c**, lower panel). The result confirms our structural observation that NME1 engages CoA and ADP by distinct interaction modes.

### The 3’ phosphate of CoA is a critical binding epitope

NME1 recognizes the 3’ phosphate group that distinguishes CoA from ADP with high specificity via five direct and two water-mediated H bonds (**Fig. 2a** and **Fig. S6a**). Although the nucleotide moiety of a CoA molecule lacking this 3’ phosphate, dephospho-CoA, has the same structure as ADP, dephospho-CoA is predicted to bind NME1 poorly because its pantoyl moiety would sterically hinder the adjacent β-phosphate from becoming buried in the active site, thereby prohibiting the canonical ADP binding mode. Indeed, NME1 incubated with dephospho-CoA yielded a native MS spectrum indistinguishable from that of the unliganded protein, confirming that NME1 does not bind dephospho-CoA (**Fig. 2c**). Thus, the 3’ phosphate of CoA is a critical feature required for NME1 recognition.

Since NME1 recognizes ADP as well as the ADP-containing end of CoA, it might conceivably bind structurally related metabolites such as NAD, FAD and ADP-ribose. To verify this, we bound NME1 to CoA beads and tested metabolites for their ability to elute the protein. As expected, NME1 was efficiently released from the beads by ATP and CoA but not by dephospho-CoA (**Fig. 2d**). NAD, FAD and ADP-ribose were equally ineffective at eluting NME1, revealing that NME1 does not promiscuously bind ADP-containing metabolites (**Fig. 2d**). Like dephospho-CoA, the lack of a 3’ phosphate group prevents these metabolites from binding like CoA, while their ADP-linked moieties sterically hinder them from burying their β-phosphate in the active site like ADP. These data suggest that CoA/acyl-CoA recognition by NME1 may be modulated by cellular ADP and ATP levels but not by other common ADP-containing metabolites.

### CoA binding and histidine phosphorylation are mutually inhibitory

Although CoA binds to the NDP/NTP binding site of NME1, previous studies have reached disparate conclusions regarding the effect of CoA on the NDPK activity of NME1/2, with one study reporting competitive inhibition by CoA in a coupled enzymatic assay^14^ and another reporting no significant inhibition by CoA or short-chain acyl-CoA in a chromatography-based assay^15^. Moreover, it is unknown whether the phosphoenzyme intermediate that mediates NDPK activity is able to bind CoA. Interestingly, another metabolite, 3’-phosphoadenosine 5’-phosphosulfate (PAPS), which chemically resembles the CoA nucleotide and adopts a similar ligand binding mode (**Fig. S10a**), was reported to inhibit *Dictyostelium discoideum* NDPK (*Dd*NDPK) with 3-fold higher binding affinity than ADP^25^. Furthermore, structurally aligning CoA-bound NME1 with an NDPK phosphoenzyme intermediate^30^ closely juxtaposes the CoA 3’ and histidine phosphate groups, which would yield a strong steric and electrostatic repulsion (**Fig. S10b**). These observations predict that CoA binding should be mutually exclusive with NDPK activity and with phosphoenzyme formation.

To verify this hypothesis, we incubated NME1 with immobilized CoA in the absence or presence of ATP and detected the total and phosphorylated protein in bound and unbound fractions by immunoblotting. Whereas in the absence of ATP most NME1 remained in the bound fraction, the addition of ATP caused the release of phosphorylated NME1 from the CoA beads, confirming that CoA binds poorly to the phosphoenzyme intermediate (**Fig. 2e**). For further proof, we used native MS to analyse NME1 hexamers following incubation with either CoA, ATP or both ligands. Incubation with CoA resulted in a predominance of NME1 hexamers bound to 3 or 4 CoA ligands, consistent with a ∼60% binding site occupancy (**Fig. 2c**, bottom right and **Fig. S11a**). Co-incubation with both CoA and ATP resulted in a leftward shift of peak intensities, revealing a prevalence of NME1 hexamers bound to only 1 or 2 CoA ligands and a ∼20% binding site occupancy (**Fig. 2c**, top right and **Fig. S11b**), confirming that ATP inhibits CoA binding. Incubation with ATP did not yield any detectable nucleotide-bound hexamers, but inspection of individual peaks revealed closely spaced sub-peaks corresponding to phosphorylated hexamers (**Fig. 2c**, lower inset), analogous to the closely spaced bands seen on a native gel (**Fig. 1e**). The majority of hexamers carried 1-3 phosphate groups, consistent with the monophosphorylation of 32% of NME1 monomers (**Fig. 2c**, middle right panel and **Fig. S11c**). In contrast, following co-incubation with both ATP and CoA the phosphorylation level of the unbound NME1 hexamer was only 16%, and this level progressively decreased to 10% for hexamers bound to either 1, 2 or 3 CoA ligands, directly demonstrating that CoA inhibits histidine phosphorylation (**Fig. 2c**, top right and **Fig. S11b**).

Taken together, these data provide compelling evidence that the NDP kinase and CoA-binding functionalities of NME1 are mutually inhibitory.

### NME1/2 repress lipogenesis in the liver

We next turned to a mouse knockout (ko) model to explore the potential importance of NME1/2 for CoA/acyl-CoA utilizing pathways. Since an *Nme1/2* double knockout (ko) is embryonic lethal, we decided to work on *Nme2* ^-/-^ (ko) mice, which present only a mild phenotype^12, 31^. An initial screen of various tissues revealed that the total level of NME1/2 is dramatically reduced in the liver of *Nme2 ^-/-^* mice compared to wild-type (*Nme2 ^+/+^*) mice (**Fig. 3a**). A CoA pull-down approach confirmed that NME1/2 were highly abundant CoA-binding factors in WT mouse liver extracts and were drastically reduced in those from *Nme2^-/-^* mice (**Fig. 3b**), identifying the liver as an expedient model system for subsequent experiments. Since the liver is one of the most active sites for *de novo* lipogenesis (DNL), which depends on a continuous supply of AcCoA, we focused on the role of NME2 in this process. MS-based lipidomic analysis of liver extracts from *Nme2^+/+^* and *^-/-^* mice revealed that the disruption of *Nme2* resulted in a significant increase in the total fatty acid content (**Fig. 3c**). A detailed analysis of hepatocyte fatty acids showed significantly increased levels of C16:0, C16:1, C18:1cis and C18:2 fatty acid species in *Nme2*^-/-^ mice (**Fig. 3d**). These findings demonstrate that NME2 negatively regulates the accumulation of cellular fatty acids.

**Figure 3.**
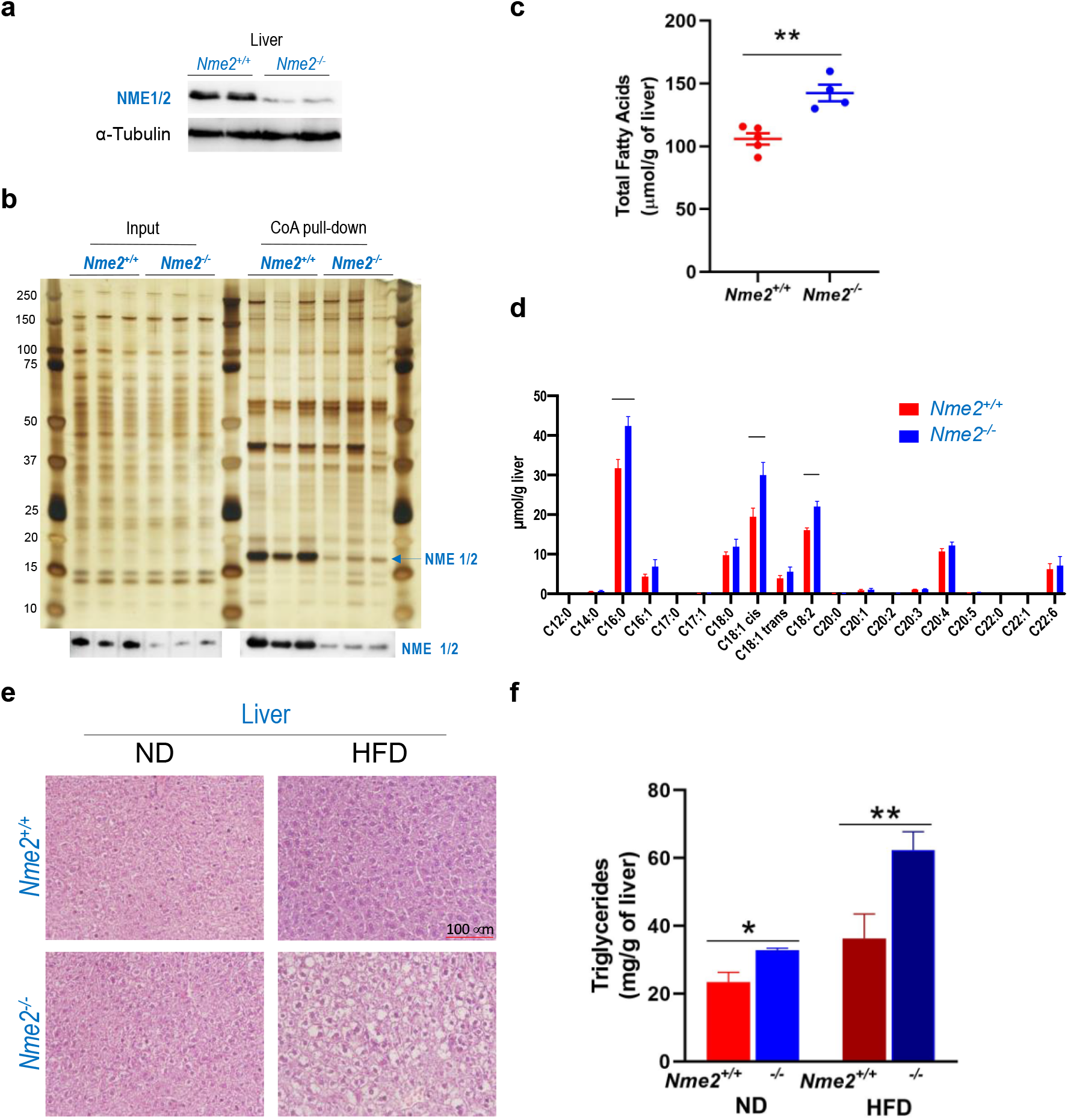
NME1/2 repress lipogenesis in the liver. **a.** Total amounts of NME1 and NME2 were visualized in the liver of *Nme2^+/+^* and *Nme2*^-/-^ mice. **b.** Liver extracts from three different *Nme2^+/+^* and *Nme2*^-/-^ mice were used in a CoA-bead pull- down experiment as described in Fig. 1b. Input (0.5 %) and CoA bound materials were visualized on a silver-stained gel and a fraction was used to show the presence of NME1/2 in the corresponding samples by immunoblotting (lower panels). **c.** Values corresponding to the total amounts of fatty acids measured in liver extracts from 5 different *Nme2^+/+^* and 4 *Nme2*^-/-^ mice are shown. \*\**p* = 0.0021. **d.** The amounts of different fatty acid species (indicated as C*x*:*y*, where *x* and *y* are the number of carbon atoms and double bonds, respectively) in liver extracts from 5 different *Nme2^+/+^* and 4 *Nme2*^-/-^ mice are shown. Error bars represent standard error of means (SEM). \*\*\*\**p*< 0.001. **_e._** *Nme2^+/+^* and *Nme2*^-/-^ mice were subject to normal diet (ND, respectively 4 *Nme2^+/+^* and 5 *Nme2*^-/-^ mice) or high fat diet (HFD, respectively 6 *Nme2^+/+^* and 7 *Nme2*^-/-^ mice). Representative images of hematoxylin and eosin (H&E)-stained paraffin-embedded liver sections from *Nme2^+/+^* and *Nme2*^-/-^ male mice after 6 weeks of ND or HFD are shown as indicated. The scale bar is 100 μm. **f.** Triglyceride concentrations were measured in liver extracts from 4 *Nme2^+/+^* and 4 *Nme2*^-/-^ mice in ND and from 10 *Nme2*^+/+^ and 12 *Nme2*^-/-^ after 6 weeks of HFD. \**p* = 0.0286 ND: *Nme2^+/+^* versus *Nme2^-/-^*; \*\**p* = 0.0071 HFD: *Nme2^+/+^* versus *Nme2^-/-^*.

### NME2 regulates a liver cell response to a high-fat diet challenge

Since a high-fat diet (HFD) is known to suppress DNL^32–35^ and we found NME2 to counteract fatty acid accumulation, we surmised a role for NME1/2 in the HFD-dependent repression of DNL. To test this idea, we fed *Nme2^+/+^* and *^-/-^* mice with a HFD or a normal diet (ND) for six weeks. Interestingly, unlike WT mice, the *Nme2^-/-^* mice exhibited clear signs of liver steatosis (**Fig. 3e**) and increased liver triglyceride levels (**Fig. 3f**) under HFD. These results suggest that, under a HFD challenge, *Nme2*^-/-^ liver cells fail to suppress the expression of genes promoting liver lipogenesis. To verify this hypothesis, we performed a transcriptomic analysis of liver cells from *Nme2 ^+/+^* and *^-/-^* mice fed normally (ND) or undergoing a HFD challenge. In *Nme2 ^+/+^* mice, the HFD challenge induced changes in the transcriptional activity of hepatocytes characterized by the marked repression and activation of 107 and 52 genes, respectively (absolute fold change ≥ 2; *p*-value ≤ 0.01) (**Fig. 4a**, *Nme2^+/+^*, ND versus HFD). *Nme2* ^-/-^ mice exhibited a strikingly different pattern, revealing that the HFD-dependent gene activation was abolished for the majority (∼70%) of genes in the latter group, while ∼20% of genes escaped the HFD-dependent transcriptional repression (**Fig. 4a**, *Nme2^-/-^*, ND versus HFD). A gene set enrichment analysis (GSEA) revealed that HFD repressed a significant number of genes involved in adipogenesis in the liver of *Nme2^+/+^* mice (**Fig. 4b**, left panel), as previously reported^35^. Remarkably, this HFD-dependent repression only occurred in the presence of NME2 (**Fig. 4b**, right panel). More specifically, in WT hepatocytes from mice under HFD, the expression of known master transcription activators of genes involved in lipid metabolism, such as PPARα, PPARγ, PPARγC1α (PGC1α) and the lipogenic genes SREBF1/CEBPα couple, was repressed, whereas in the absence of NME2, these genes either failed to be repressed or were even upregulated despite the HFD challenge (**Fig. 4c**). The continuous expression of these master transcription factors explains the inability of *Nme2*^-/-^ hepatocytes to repress lipogenic genes in response to a HFD challenge.

**Figure 4.**
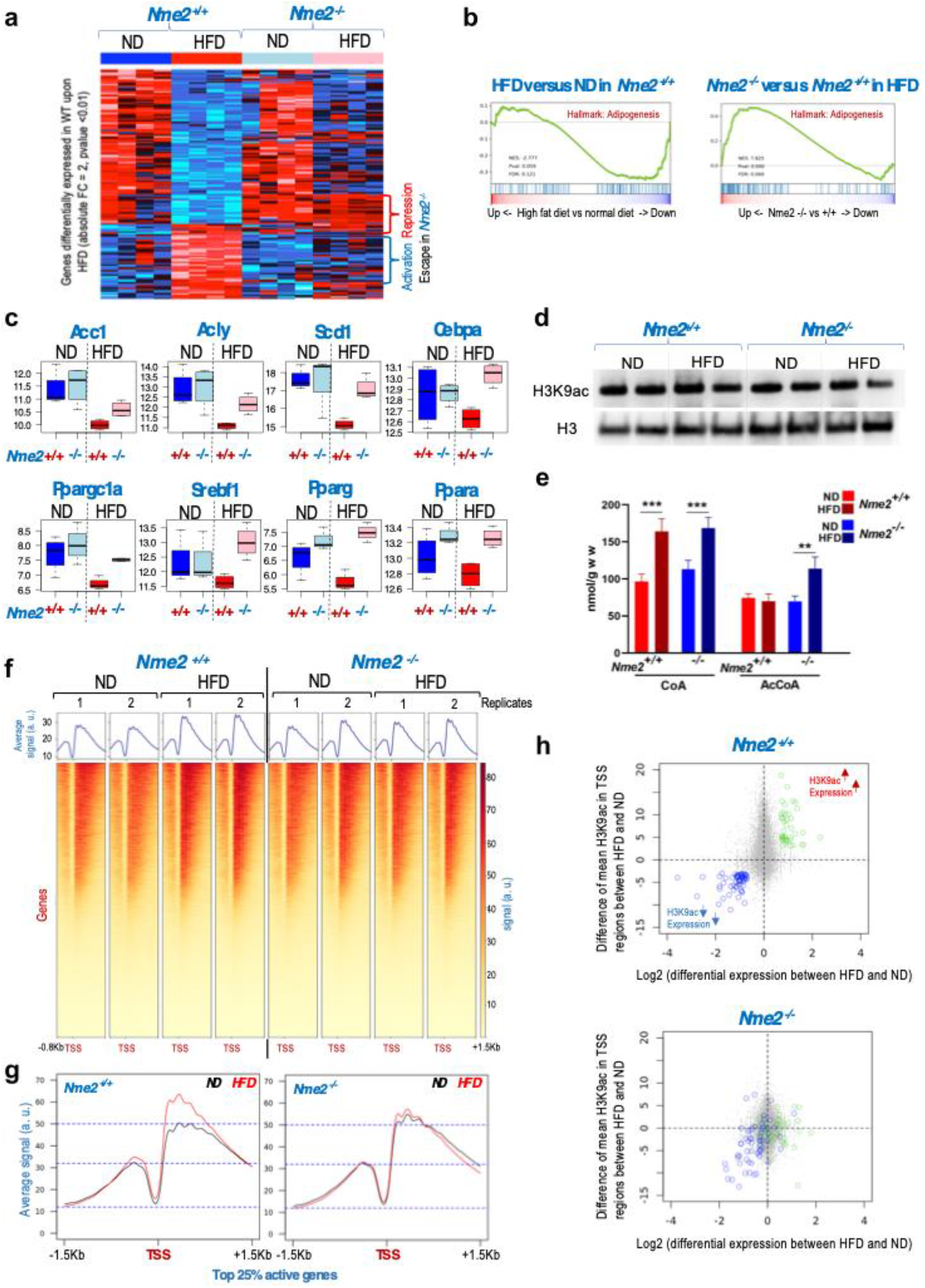
NME2 regulates the liver cell response to a high-fat diet challenge. **a.** RNAs extracted from liver of four mice of each indicated genotype kept under normal diet (ND) or fed for six weeks with a high fat diet (HFD) were subjected to RNA sequencing. The expression of genes differentially expressed between the liver of HFD and ND fed *Nme2*^+/+^ mice (with a fold change absolute value >2 and a *p*-value <0.01) is shown as a heatmap in *Nme2*^+/+^ and *Nme2^-/-^* mice for both conditions as indicated. Hierarchical clustering enables the identification of genes that escape HFD-dependent repression or activation in *Nme2^-/-^* mice (indicated on the right side of the heatmap by red and blue brackets, respectively). **b.** GSEA plots showing a downregulation of the geneset corresponding to genes involved in adipogenesis in the hepatocytes of *Nme2*^+/+^ mice liver treated with HFD for 6 weeks compared with ND (reference, left panel). The same geneset is significantly enriched in hepatocytes of *Nme2^-/-^* mice compared to *Nme2^+/+^* mice (reference) under HFD (right panel), showing that the HFD-induced repression of this geneset is abolished in *Nme2^-/-^* hepatocytes. **c.** Box plots representing the distribution of the expression levels of the indicated individual genes in liver from *Nme2*^+/+^or *Nme2^-/-^* mice under normal (ND) or high fat diet (HFD) are shown. The values correspond to the normalized RNAseq read counts. **d.** The levels of H3K9 acetylation and histone H3 in the liver extracts from the mouse treated as above were visualized by immunoblots using the corresponding antibodies. Two samples per condition were collected from independent mice. **e.** The cellular concentrations of CoA and AcCoA were respectively measured in liver extracts from *Nme2*^+/+^ and *Nme2^-/-^* mice. For each measurement, livers from independent mice were used as follows: *Nme2*^+/+^ ND, n = 9; *Nme2*^+/+^ HFD, n = 4; *Nme2^-/-^* ND, n = 9; *Nme2^-/-^* HFD, n = 5. The graphs show the average of the concentration values (in nmol/g of liver) and ± SEM. CoA: *** *p* = 0.003 for HFD vs ND in *Nme2^+/+^* and *p* = 0.007 for HFD vs ND *Nme2^-/-^*. AcCoA: ** *p* = 0.002 for HFD vs ND in *Nme2^-/-^*. **f.** Nuclei from the livers of 2 independent mice for each condition (as indicated) were purified and extensively digested with Mnase to release mono-nucleosomes, which were then immunoprecipitated using an anti-H3K9ac antibody. The DNA associated with the immunoprecipitated nucleosomes was sequenced and the normalized read counts at the corresponding gene TSS (-/+ 5 Kb) are visualized in this heatmap. The normalization procedure assumed that the total levels of H3K9ac were similar in all samples as demonstrated by the immunoblots shown in Fig. 4d. The TSS regions are ranked as a function of the mean signal value across all conditions, from the highest to the lowest. **g.** The mean value of two independent H3K9ac ChIP read counts from HFD (red) and ND (black) in *Nme2^+/+^* and *Nme2^-/-^* mice were plotted over +/-1.5 Kb centered on the TSS regions corresponding to the 25% top most highly expressed genes (according to the transcriptomic data of the liver samples from *Nme2^+/+^* mice in normal diet, n = 3500 TSS). **h.** The difference of the mean H3K9ac ChIP-seq values between HFD and ND was calculated for each of the TSS-0.8+1.5Kb regions (y-axis) and plotted against the differential expression values (log2 of the fold change) of the corresponding genes between the same conditions (y-axis) respectively for *Nme2^+/+^* (upper panel) or *Nme2^-/-^* (lower panel) mice. Considering the *Nme2^+/+^* liver samples (upper panel), two groups of TSS/genes were selected, either presenting both an enhanced expression and an increased TSS H3K9ac level between HFD and ND (green dots) or showing a reduced expression and a decreased H3K9ac level between HFD and ND (blue dots). The same genes are visualized on the plot corresponding to the *Nme2^-/-^* liver in the lower panel (the genes symbolized by green or blue dots are the same in both panels).

The GSEA also revealed specific genes that were upregulated in response to a HFD challenge in a NME2-dependent manner. These genes are mostly involved in the IL6/Jak/Stat3, interferon α and γ, and TNFα pathways (**Fig. S12**). Interestingly, all these gene expression pathways have been previously shown to play a critical protective role during HFD-induced liver injury and liver regeneration^36^. Altogether, the above transcriptomic analyses suggest that NME2 could be a general regulator of a protective transcriptional response of liver cells to a HFD challenge that acts by suppressing DNL and activating a protective cytokine response.

In order to exclude a role for AMPK (known to phosphorylate and inactivate ACC1) in the regulation of lipogenesis by NME1/2, we used immunoblotting to monitor the overall expression and phosphorylation of ACC1 and AMPK proteins (**Fig. S13a**). As expected, the variations of ACC1 mirrored that of its encoding mRNA under the different conditions tested (Compare **Fig. S13a** and **Fig. 4c**, ACC1 panels). Importantly, the level of inactivating ACC1 phosphorylation by AMPK remained unchanged between *Nme2*^-/-^ and *^+/+^* livers, consistent with unchanged levels of AMPK activation (P-AMPK/AMPK ratio) and of cellular ATP, ADP and AMP under the different tested conditions (**Fig. S13b**). Thus, AMPK signalling is not involved in ACC1 regulation or in the increased DNL under these conditions. Overall, these data demonstrate that NME2 directly controls the liver cell response to HFD.

### NME2-dependent redistribution of H3K9ac at gene transcriptional start sites (TSSs)

We next investigated whether the transcriptional regulation observed in liver under HFD involved an NME2-dependent control of histone acetylation. We focused on H3K9ac as a histone mark known to be associated with active gene transcription and responsive to a metabolic change^37, 38^. **Fig. 4d** shows that a HFD challenge or *Nme2* ^-*/-*^ did not change the global level of H3K9ac in hepatocytes, which was comparable across all tested conditions. Measurement of the cellular CoA and AcCoA concentrations showed that, although the HFD challenge increased the amount of CoA in both WT and ko cells, it did not significantly alter the amount of AcCoA in WT cells (**Fig. 4e**). Interestingly, in the absence of NME2, a HFD challenge led to a significant (1.65- fold relative to ND) increase in AcCoA, which could contribute to the observed increase in fatty acid synthesis (**Fig. 4e**).

Using a ChIP-seq approach we then investigated whether NME2 was involved in controlling the distribution of H3K9ac at gene TSSs, which would explain the transcriptional reprograming of HFD-dependent genes. **Figures 4f** and **g** show that, in *Nme2 ^+/+^* liver cells, the HFD challenge led to an increase of H3K9ac at gene TSSs, which was especially pronounced at highly active genes. The enhanced acetylation occurred on nucleosomes (primarily at positions +1, +2 and +3) downstream of the TSS of active genes but not on upstream nucleosomes (**Fig. 4g**, left panel). Strikingly, this redistribution of H3K9ac did not occur in *Nme2* ^-/-^ liver cells (**Fig. 4g**, right panel).

Enhanced H3K9ac downstream of TSSs is known to promote RNA pol II pause release^39^ and thereby enhance transcription. To correlate the redistribution of H3K9ac at TSS-associated nucleosomes with transcriptional activity^40^, we highlighted genes whose acetylation and transcription levels either both increased or both decreased under HFD compared to ND in the liver of *Nme2 ^+/+^* mice (**Fig. 4h**, upper panel, green and blue dots respectively). Thus, the redistribution of H3K9ac on the TSS regions of these genes is associated with the transcriptional response to HFD in *Nme2 ^+/+^* hepatocytes. In *Nme2* ^-/-^ hepatocytes, the HFD-dependant H3K9ac redistribution and transcriptional response to HFD was lost for many of these genes, including nearly all those which normally showed a combined increase in expression and TSS-associated H3K9ac (**Fig. 4h**, lower panel, green dots). NME2 seems required for the combined decrease in expression and TSS-associated H3K9ac for only a subset of genes under HFD (**Fig. 4h**, lower panel, blue dots). However, these genes include the critical regulators of lipogenic genes (**Fig. 4c**).

### AcCoA is critical for the NME1/2-dependent control of lipogenic gene repression

To assess whether NME1/2 also controlled the expression of lipogenic genes in human liver cells, we used the established hepatocellular carcinoma cell line, HepG2. In these cells, *Nme1/2* knock-down led to a remarkable accumulation of ACC1 (**Fig. 5a**, si*Nme1*), suggesting a similar regulatory effect of NME1/2 on DNL as in mouse liver cells (i.e., activation of DNL in the absence of NME2). The observed ACC1 accumulation suggests that DNL in WT HepG2 cells is significantly repressed in an NME1/2-dependent manner and provides a useful readout of DNL activation. Indeed, as shown in **Fig. 4c** and **Fig. S13a**, active DNL under ND is associated with enhanced expression of ACC1 compared to repressed DNL (under HFD). Upregulation of liver ACC1 following the activation of DNL has also been reported in other contexts such as high-carbohydrate/low fat diet^41^ and acetate- or fructose-induced DNL^42, 43^

**Figure 5.**
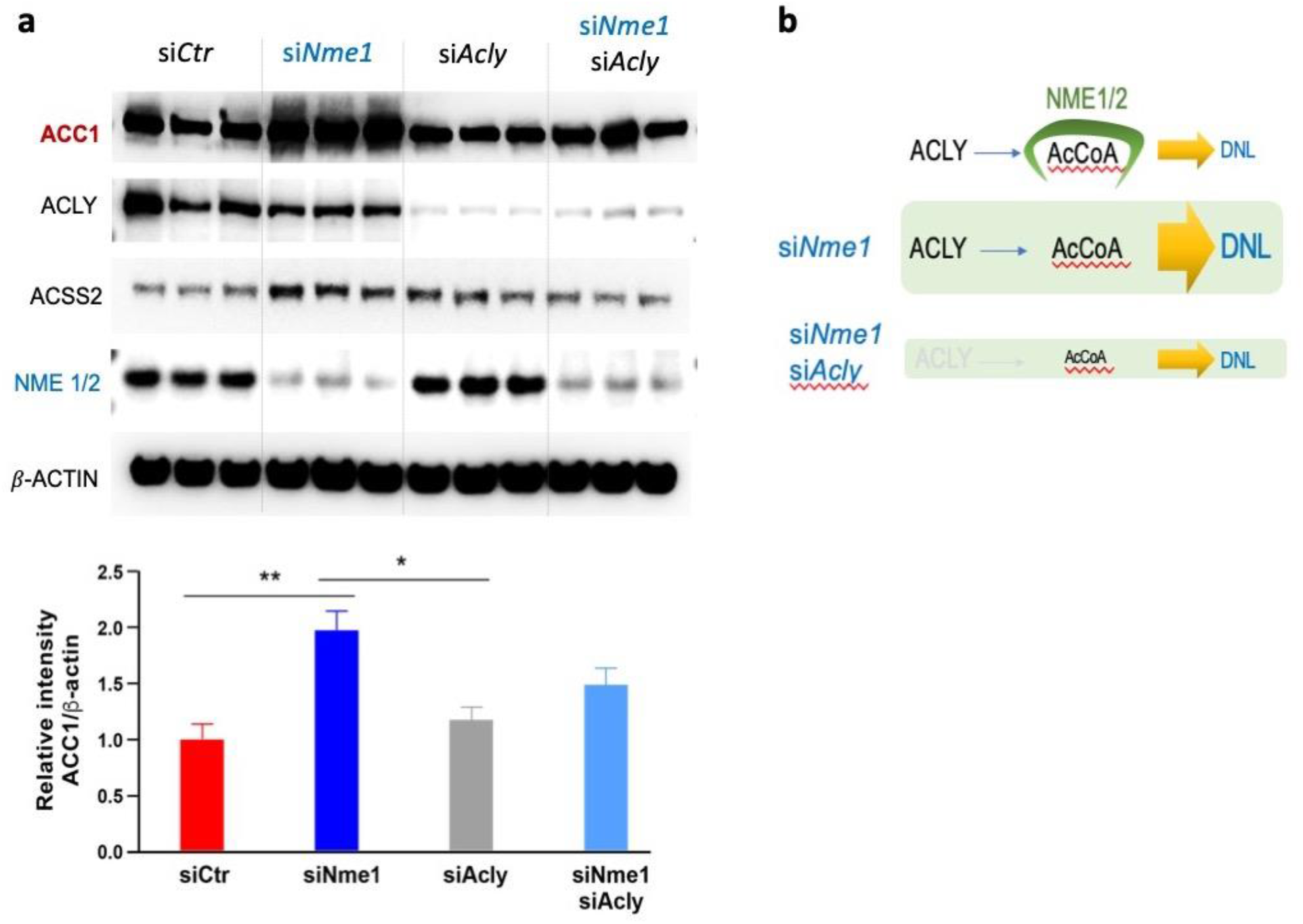
CoA/AcCoA binding by NME1/2 is critical for lipogenic gene repression. **a.** *Nme1* or *Acly* or both were knocked down in HepG2 hepatocellular carcinoma cells and the levels of the indicated proteins were visualized by immunodetection using the corresponding antibodies. The bar plot underneath shows, for each condition, the average of relative intensity ACC1/β-actin values and corresponding ±SEM obtained from three independent cell cultures. Significance of one-way ANOVA tests: \*\**p*=0.0059 for siCTR versus siNme1, and \**p*=0.0180 for si*Nme1* versus si*Acly*. **b.** Scheme illustrating a possible interpretation of the observed data. In wild-type cells NME1/2 sequester AcCoA from DNL. NME1/2 knock-down makes AcCoA freely available to ACC1 thereby enhancing DNL. The NME1/2 knockdown effect on DNL is abolished by reducing AcCoA production following the knock-down of ACLY.

ACC enzymes recognize the same nucleotide moiety of CoA as does NME1^44^, implying that AcCoA binding by ACC1 and NME1 is mutually exclusive. Consequently, an attractive scenario would be that NME1/2 restricts the availability of AcCoA to the DNL machinery, whereas NME1/2 depletion would make AcCoA more available to fuel DNL. One prediction of this model is that a decrease in AcCoA production should abolish the DNL activation observed upon *Nme1/2* knock- down. Since the ATP citrate lyase (ACLY)-produced AcCoA pool constitutes an important source of cytoplasmic AcCoA, we performed a double knock-down of *Acly* and *Nme1* and monitored the accumulation of ACC1. **Fig. 5a** shows that ACLY downregulation abolished the accumulation of ACC1 induced by the *Nme1* knock-down, strongly supporting the hypothesis that NME1/2 restricts AcCoA availability to DNL (**Fig. 5b**). This hypothesis is also in line with an enhanced AcCoA synthesis in *Nme2* ko liver cells (**Fig. 4e**), which could reflect a better accessibility of CoA to AcCoA synthesizing enzymes, ACLY and ACSS2. Admittedly, the ideal experiment to test the above hypothesis would have involved using an NME1 mutant specifically compromised for binding CoA while retaining full kinase activity. Unfortunately, we were unable to perform such an experiment since the kinase activity of our NME1 T94D mutant was significantly reduced (**Fig. S9d**). Therefore, manipulating AcCoA production in the context of wild-type or knock-down NME1/2 cells emerged as the best alternative approach for confirming the role of NME1/2 in restricting the availability of AcCoA to fuel ACC1 activity.

Taken together, this work demonstrates that NME1/2 play a critical role in regulating DNL in liver cells, more specifically under HFD conditions, and suggests that this role is potentially achieved by directly controlling the ACC1–AcCoA axis.

## DISCUSSION

In this study, we showed that NME1 binds AcCoA and CoA with similar affinity via a binding mode distinct from that of canonical ADP and ATP substrates and critically dependent on the CoA 3’ phosphate group. This plasticity of ligand recognition is primarily achieved through a versatile array of structural waters and the lack of base-constraining H-bonding groups in the nucleotide binding site. These structural features make NME1/2 uniquely adapted to recognize ATP and CoA through the same binding site and thereby monitor the cellular balance between these key metabolites. This observation, together with the prevalence of NME1/2 over other cellular CoA- binding factors in our CoA-pull downs, led us to consider the role played by NME1/2 in lipogenesis, a pathway controlled by the cellular energy status and a major consumer of AcCoA, the most abundant cellular CoA derivative. The absence of a compensatory NME1/2 accumulation in the liver of *Nme2* ^-/-^ mice identified this organ, a major site of lipogenesis, as a good model for exploring the role of NME1/2 in DNL.

A lipidomic analysis of the liver from *Nme2 ^+/+^* and *^-/-^* mice revealed a repressive role for NME1/2 in lipogenesis. Remarkably, we observed that NME2 depletion abolishes the well-documented ability of hepatocytes to repress DNL under HFD. Exploring the underlying mechanism revealed that, whereas a HFD challenge normally represses the expression of major lipogenic genes, as previously reported^35^, in *Nme2 ^-/-^* hepatocytes under the same HFD conditions, these genes remain active. More specifically, we observed that, while HFD ordinarily represses the expression of master transcription factors involved in lipid metabolism such as PPARα, PPARγ, PPARγC1α (PGC1α) and SREBF1/CEBPα, in the absence of NME2 the expression of these factors persists. In agreement with the role of these factors as transcription drivers, we observed that a series of other genes involved in adipogenesis also escape repression under HFD in *Nme2 ^-/-^* liver cells. HFD also induces the expression of multiple genes known to play a protective role during liver regeneration, including those involved in the IL6/Jak/Stat3, interferon α and γ, and TNFα pathways^36^. These transcriptomic analyses therefore suggest that NME1/2 could be general regulators of the liver response to stressful assaults.

A genome-wide mapping of H3K9ac in *Nme2^+/+^* and *Nme2^-/-^* liver cells from mice under normal diet or HFD challenge also revealed the occurrence of an HFD-induced increase of H3K9ac at active gene TSSs. Interestingly, this specific accumulation of H3K9ac was completely abolished in the absence of NME2. Based on the known competition between DNL and histone acetylation^45–47^, a possible explanation could be that the NME1/2-dependent down-regulation of DNL under HFD makes the AcCoA pool more available for histone acetylation. In the absence of NME1/2 and the inability of HFD to repress DNL, AcCoA could continue to be essentially used by DNL, explaining why no change in histone acetylation was observed in *Nme2* ko liver cells.

All these findings raise the intriguing possibility that the ability of NME1/2 to bind CoA/AcCoA may play a major role in the control of DNL. Notably, NME1/2 are sufficiently abundant to have a significant buffering effect on the free CoA/AcCoA levels in the cytosol. Our measurements indicate that the combined concentration of CoA and AcCoA in mouse liver cells remains below 200 μM, consistent with previous estimates in rat liver cells (90-100 μM)^48, 49^ and mammalian cells in general (20-140 μM)^50^. However, most of these molecules (90-95%) will be in the mitochondria^49, 51^ and an additional fraction will be bound by various carriers and enzymes, and so the available cytosolic concentration is likely to be in the 1-20 μM range. NME proteins are among the 100 most abundant cellular proteins from *E. coli*^52^ to human^53^. Although the NME1/2 concentration in mouse liver cells is unknown, a study of mouse fibroblasts reported an NME1 copy number of >21 million molecules per cell^54^. Assuming a cell volume of 2000 μm^3^ (ref.^55^), this implies an NME1 concentration of ∼18 μM, closely matching the *K*_D_ value we measured for CoA binding. The fact that both values are commensurate with the cytosolic CoA/AcCoA concentration means that a significant fraction (up to 40-50%) of these molecules are potentially bound by NME1/2, in line with the observed prevalence of NME1/2 over other cellular CoA/AcCoA-binding factors in our CoA pull-downs.

Interestingly, NME1/2 are known to associate with DNA and with numerous transcription factors and transcriptional regulators^56, 57^. The DNA-binding surface of NME2 has been mapped to residues near the dyad axis on the outer equatorial surface of the hexamer, far from the nucleotide-binding site^58^, suggesting that NME1/2 could feasibly bind AcCoA and DNA simultaneously. These interactions would place NME1/2 in the vicinity of chromatin-associated HATs, where they could promote histone acetylation by facilitating the delivery of AcCoA to the HAT active site. Indeed, many HATs have *K*_D_ and *K*_m_ values for AcCoA (0.5 - 3 μM)^59–63^ lower than the *K*_D_ we measured for NME1 (18 μM), implying a favourable affinity gradient from NME1/2 to HATs.

While previous studies showed that the NDPK activity of NME1/2 is negatively regulated by CoA or LCFA-CoA^14, 15^, we additionally established that the converse is true, i.e., that CoA binding is inhibited by NDPK activity, specifically, by the phosphoenzyme intermediate that lies along the reaction pathway. Since the degree of histidine phosphorylation varies with ATP concentration, this means that the NME1/2:AcCoA interaction is sensitive to the energy status of the cell, raising the possibility that NME1/2 may dynamically sequester AcCoA molecules in an ATP-dependent manner. This mechanism would provide an additional level of control of lipogenesis that would synergize with the well-known AMPK-dependent energy sensing mechanism. Indeed, high cellular ATP concentrations would not only prevent the inactivating phosphorylation of ACC1 by AMPK, but, by inhibiting sequestration by NME1/2, would also make more AcCoA available to ACC1, leading to enhanced DNL and a consequent increase of ACC1 accumulation^41–43^. One potential objection to this idea is that the levels of ATP in the cell (usually in the low mM range^64^) significantly exceed those of cytosolic AcCoA and hence should outcompete AcCoA for binding to NME1. However, our native MS experiments showed that even in the presence of millimolar ATP, only a subset of NME1 monomers within each hexamer was phosphorylated and that no stable ADP- or ATP-bound species were detected, indicating that the remaining monomers were free to bind CoA.

In conclusion, the work reported here highlights the function of NME1/2 as a major repressor of DNL in hepatocytes and responsible for a liver protective gene expression program. In agreement with this finding, a previous study investigating genetic factors controlling liver injury susceptibility identified NME1/2 as important determinants in protecting the liver against an injury-inducing treatment^65^. Therefore, NME1/2 may be a general regulator of a protective liver response in various pathophysiological contexts.

## ACKNOWLEDGEMENTS

This work was supported by the “Université Grenoble Alpes” ANR-15-IDEX-02 SYMER (CP, US, SK) and LIFE (SK) programs, by the Agence Nationale de la Recherche (ANR) Episperm4 (ANR-19-CE12- 0014) program (SK, CP), by Plan Cancer Pitcher and from MSD Avenir ERICAN programs (SK), as well as by the Cancer ITMO (Multi-Organisation Thematic Institute) of the French Alliance for Life Sciences and Health (AVIESAN) MIC program (SK and SR), and the Institut Universitaire de France (US). The proteomic experiments were partially supported by Agence Nationale de la Recherche under projects ProFI (Proteomics French Infrastructure, ANR-10-INBS-08) and GRAL, a program from the Chemistry Biology Health (CBH) Graduate School of University Grenoble Alpes (ANR-17- EURE-0003). High throughput sequencing was performed at the TGML Platform, supported by grants from Inserm, GIS IbiSA, Aix-Marseille Université, and ANR-10-INBS-0009-10. Computations presented in this paper were performed using the GRICAD infrastructure (https://gricad.univ-grenoble-alpes.fr), which is supported by Grenoble research communities. Structural, biophysical and native MS studies used the platforms of the Grenoble Instruct-ERIC center (ISBG; UMS 3518 CNRS-CEA-UGA-EMBL) within the Grenoble Partnership for Structural Biology (PSB), supported by FRISBI (ANR-10-INBS-05-02) and GRAL, financed within the University Grenoble Alpes graduate school (Ecoles Universitaires de Recherche) CBH-EUR-GS (ANR-17-EURE-0003). We thank Caroline Mas for assistance with ITC experiments. We acknowledge the European Synchrotron Radiation Facility (ESRF) for provision of synchrotron radiation facilities and thank staff of the ESRF and European Molecular Biology Laboratory (EMBL) for assistance at beamline ID30A-1. The IBS acknowledges integration into the Interdisciplinary Research Institute of Grenoble (IRIG, CEA). CYB and YYB are supported by Agence Nationale de la Recherche, France (Project ApicoLipiAdapt grant ANR-21-CE44-0010), the Fondation pour la Recherche Médicale (FRM EQU202103012700), Laboratoire d’Excellence Parafrap, France (grant ANR-11-LABX-0024), LIA-IRP CNRS Program (Apicolipid project), the Université Grenoble Alpes (IDEX ISP Apicolipid) and Région Auvergne Rhone-Alpes for the lipidomics analyses platform (Grant IRICE Project GEMELI).

## AUTHOR CONTRIBUTIONS

D.I. conceived and performed most of the reported experiments, conceived and worked with mouse models, cell lines and worked with recombinant proteins for biochemical tests and structural biology. I.G-S. performed the crystallographic work and structural analysis. Y.C. and A.A. performed proteomic analyses and interpretation and presentation of the corresponding data. Y.Y-B. performed lipidomic analyses and data presentation. N.Z. and M.T-S. and K.C. performed measurements of metabolite concentration, NME kinase activity assays and AMPK immunoblots. Z.M-J. helped with triglyceride measurements and mouse liver data interpretation. T.D. helped with liver data interpretation and significance with regard to human liver pathologies. F.B. and S.B. helped with mouse model acquisition, crossing, genotyping and breeding. F.B. also conceived the high fat diet experiment. L.S. and E.B.E. performed the LC-ESI-TOF and native MS experiments and data analysis. A.H. helped with monitoring histone modifications in cell models (data not presented). F.C. and D. P. generated transcriptomic and ChIP-seq raw data and data presentation.

S.R. coordinated bioinformatic data generation and interpretation and also generated figures (heatmaps and GSEA). L.B. sequenced RNA and ChIPed materials. A-L.V. helped with cell culture and cell line generation. C.B. coordinated the lipidomic data generation and data interpretation. All authors have read the manuscript and agreed with its content. U.S. coordinated and supervised metabolite measurement, data generation and interpretation and acted as an expert in the field of NME biology. C.P. conceived and coordinated the structural work, contributed to data interpretation and wrote the manuscript. S.K. conceived the whole study, coordinated functional studies, contributed to data interpretation and wrote the manuscript.

## DECLARATION OF INTERESTS

All the authors declare no conflict of interest.

## Supplementary Information

**Table S1.** MS-based characterization of CoA-binding proteins.

**Table S2.**
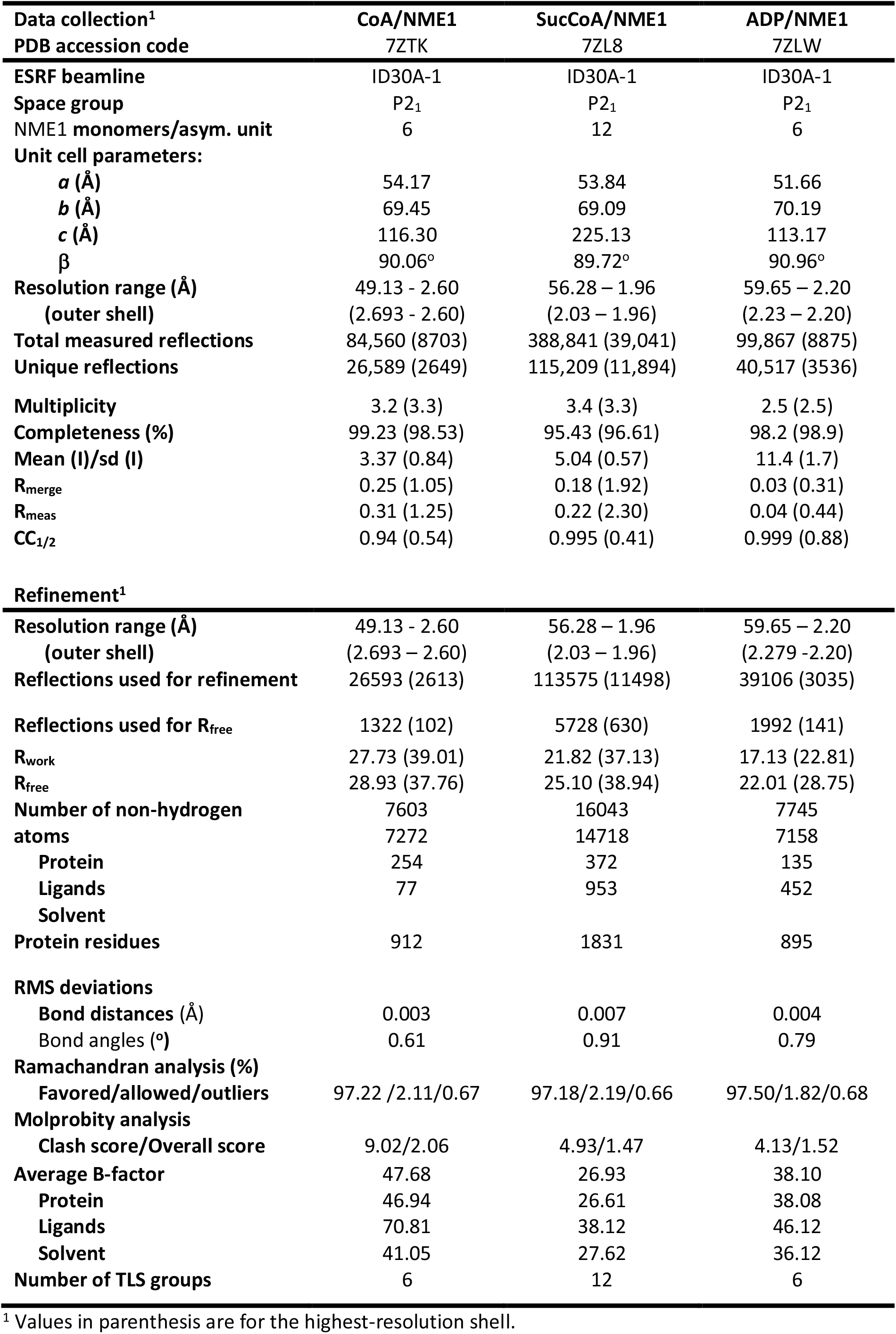
Crystallographic data collection and refinement statistics.

**Table S3.**
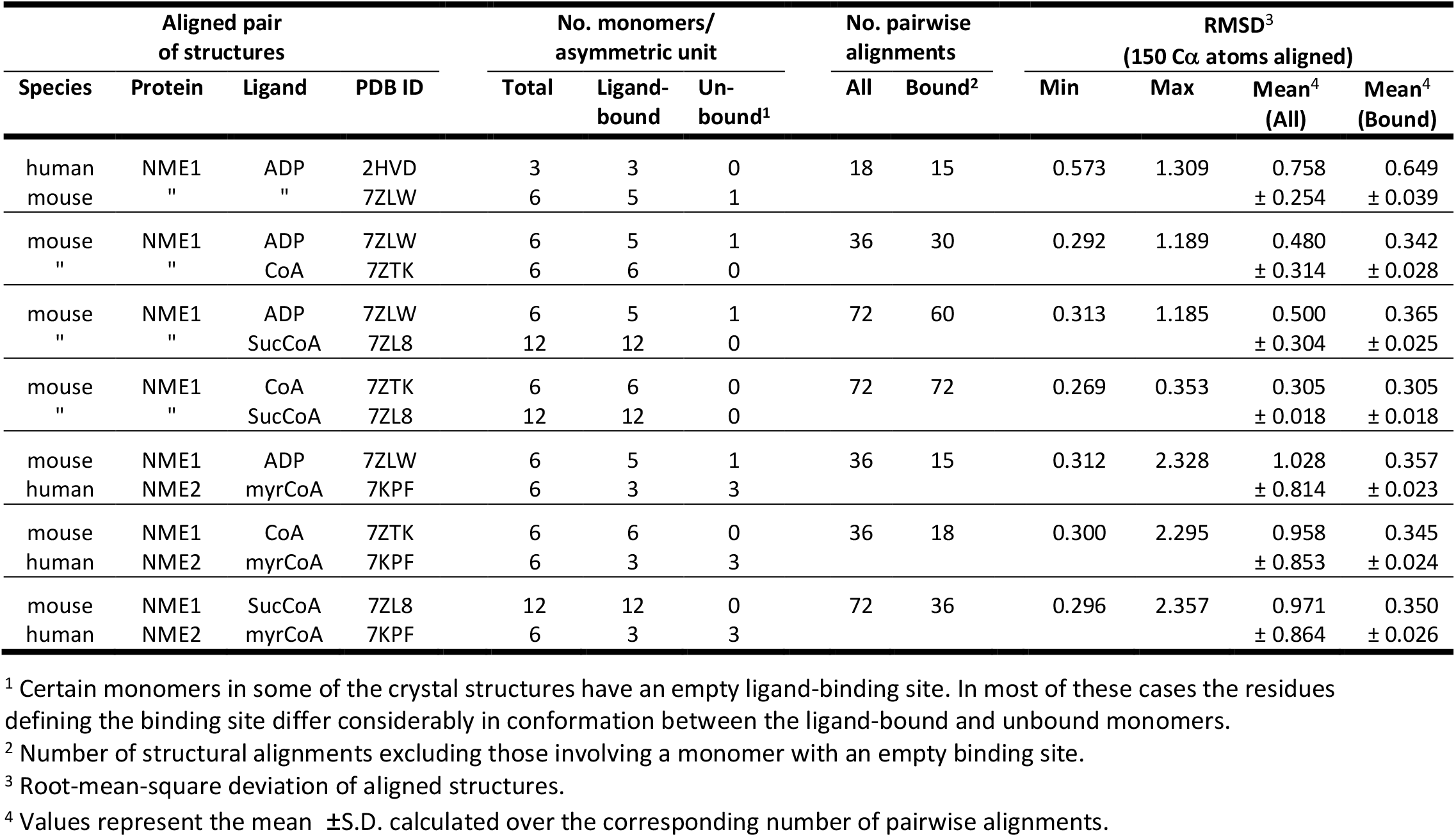
RMS differences between ligand-bound NME1/2 structures.

| Aligned pair of structures | | | | No. monomers/asymmetric unit | | | No. pairwise alignments | | RMSD<sup>3</sup><br>(150 Cα atoms aligned) | | | |
| --- | --- | --- | --- | --- | --- | --- | --- | --- | --- | --- | --- | --- |
| Species | Protein | Ligand | PDB ID | Total | Ligand-bound | Un-bound<sup>1</sup> | All | Bound<sup>2</sup> | Min | Max | Mean<sup>4</sup><br>(All) | Mean<sup>4</sup><br>(Bound) |
| human | NME1 | ADP | 2HVD | 3 | 3 | 0 | 18 | 15 | 0.573 | 1.309 | 0.758 | 0.649 |
| mouse | " | " | 7ZLW | 6 | 5 | 1 | | | | | $\pm 0.254$ | $\pm 0.039$ |
| mouse | NME1 | ADP | 7ZLW | 6 | 5 | 1 | 36 | 30 | 0.292 | 1.189 | 0.480 | 0.342 |
| " | " | CoA | 7ZTK | 6 | 6 | 0 | | | | | $\pm 0.314$ | $\pm 0.028$ |
| mouse | NME1 | ADP | 7ZLW | 6 | 5 | 1 | 72 | 60 | 0.313 | 1.185 | 0.500 | 0.365 |
| " | " | SucCoA | 7ZL8 | 12 | 12 | 0 | | | | | $\pm 0.304$ | $\pm 0.025$ |
| mouse | NME1 | CoA | 7ZTK | 6 | 6 | 0 | 72 | 72 | 0.269 | 0.353 | 0.305 | 0.305 |
| " | " | SucCoA | 7ZL8 | 12 | 12 | 0 | | | | | $\pm 0.018$ | $\pm 0.018$ |
| mouse | NME1 | ADP | 7ZLW | 6 | 5 | 1 | 36 | 15 | 0.312 | 2.328 | 1.028 | 0.357 |
| human | NME2 | myrCoA | 7KPF | 6 | 3 | 3 | | | | | $\pm 0.814$ | $\pm 0.023$ |
| mouse | NME1 | CoA | 7ZTK | 6 | 6 | 0 | 36 | 18 | 0.300 | 2.295 | 0.958 | 0.345 |
| human | NME2 | myrCoA | 7KPF | 6 | 3 | 3 | | | | | $\pm 0.853$ | $\pm 0.024$ |
| mouse | NME1 | SucCoA | 7ZL8 | 12 | 12 | 0 | 72 | 36 | 0.296 | 2.357 | 0.971 | 0.350 |
| human | NME2 | myrCoA | 7KPF | 6 | 3 | 3 | | | | | $\pm 0.864$ | $\pm 0.026$ |

<sup>1</sup> Certain monomers in some of the crystal structures have an empty ligand-binding site. In most of these cases the residues defining the binding site differ considerably in conformation between the ligand-bound and unbound monomers.
<sup>2</sup> Number of structural alignments excluding those involving a monomer with an empty binding site.
<sup>3</sup> Root-mean-square deviation of aligned structures.
<sup>4</sup> Values represent the mean ± S.D. calculated over the corresponding number of pairwise alignments.

**Figure S1.**
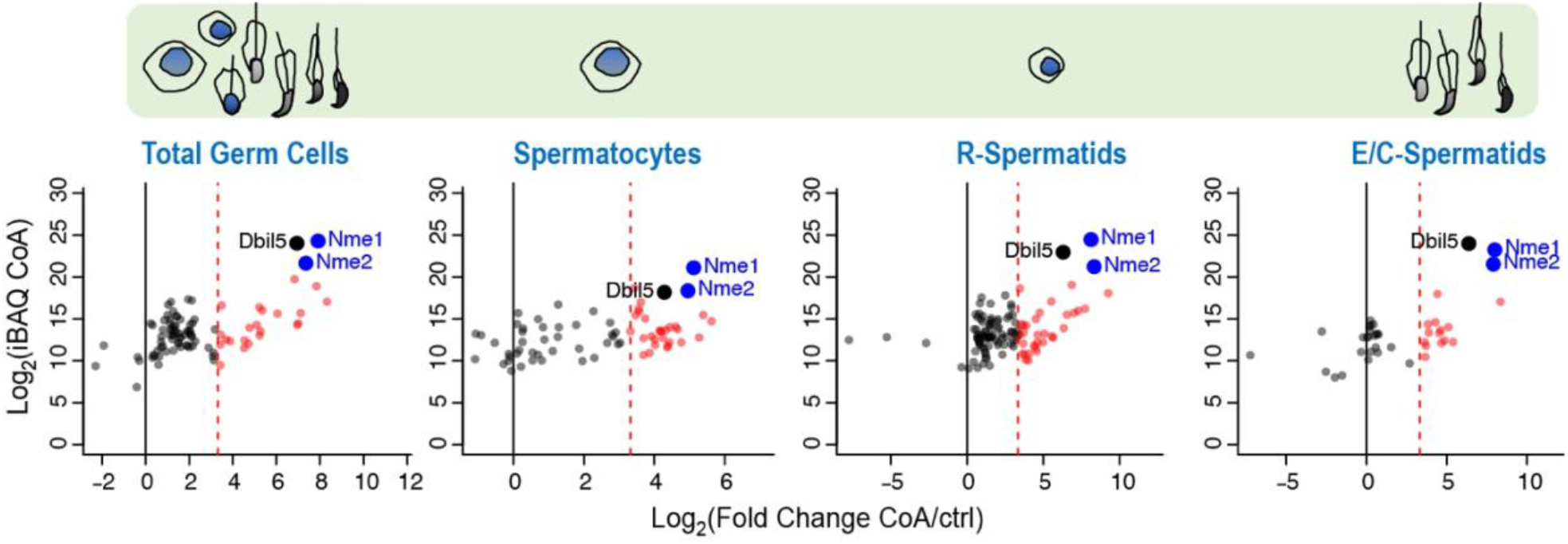
NME1/2 are major CoA-binding factors in mouse spermatogenic cells. CoA-pull downs were performed in the presence of 500 mM KCl on extracts from total germ cells, pachytene spermatocytes, round spermatids, and elongating and condensing spermatids.

**Figure S2.**
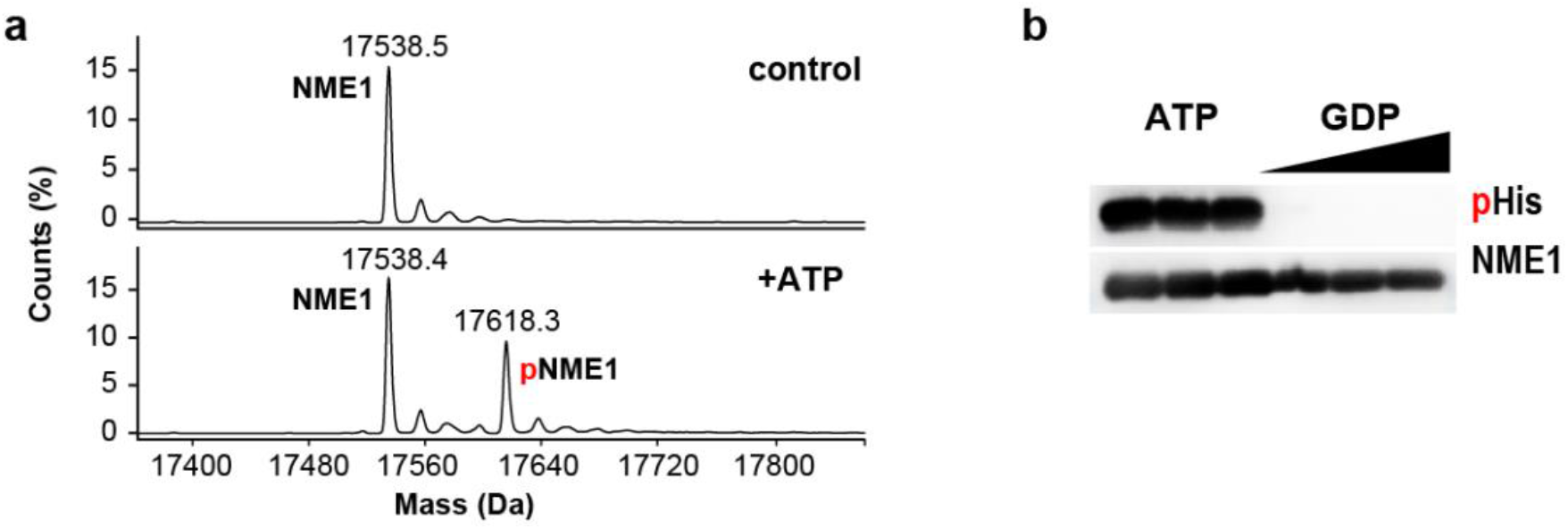
Recombinant NME1 is phosphorylated by ATP and dephosphorylated by GDP. **a.** MS analysis of NME1 incubated without (top) or with (bottom) ATP. Under the experimental conditions used (5.7 μM NME1, 100 μM ATP) approximately 40% of NME1 appears to be phosphorylated. **b.** NDPK activity confirmed by an immunoblot. An anti-phosphohistidine antibody detects phosphorylated recombinant NME1 (0.23 mM incubated with 0.5 mM ATP in triplicate). The phosphohistidine signal is lost after the incubation of phosphorylated NME1 (0.028 mM) with GDP (0.1, 0.2 and 0.4 mM).

**Figure S3.**
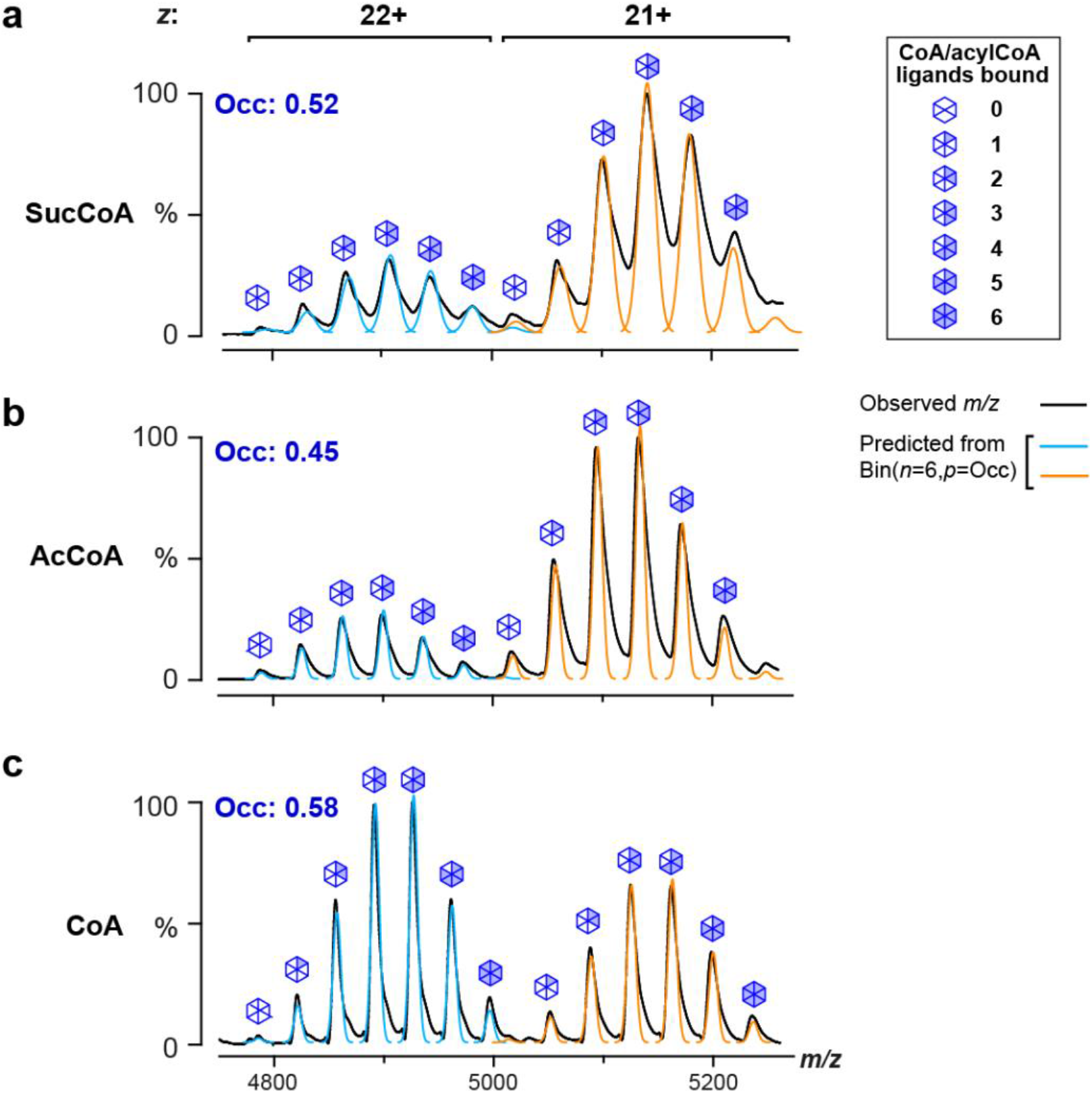
Native MS spectra reveal a binomial distribution of CoA/acyl-CoA-bound hexameric NME1 species. Spectra shown are the same as in **Fig. 1h**. Peaks in the spectra were modelled as a simple Gaussian function whose amplitude was scaled by the values given by the binomial distribution, Bin(*N*,*p*), where *N* is the number of binding sites on the NME1 hexamer (*N*=6) and *p* is the fitted probability of a single site being occupied by a ligand. The fraction of NME1 hexamers bound to *r* ligands (*r* = 0 to 6) is given by C(6,*r*)*p^r^*(1-*p*)^1-*r*^ where C(6,r)= 6!/[r!(6-r)!]. Recorded spectra are shown in black and fitted curves for the 22+ and 21+ charge states are in cyan and orange, respectively. The good agreement between the recorded spectra and the fitted curves confirms that CoA/acylCoA ligands bind independently to the six binding sites on each NME1 hexamer, allowing one to estimate the fraction (*Occ*, equal to *p*) of NME1 monomers bound to each ligand, which for these experiments ranged between 0.45 and 0.58.

**Figure S4.**
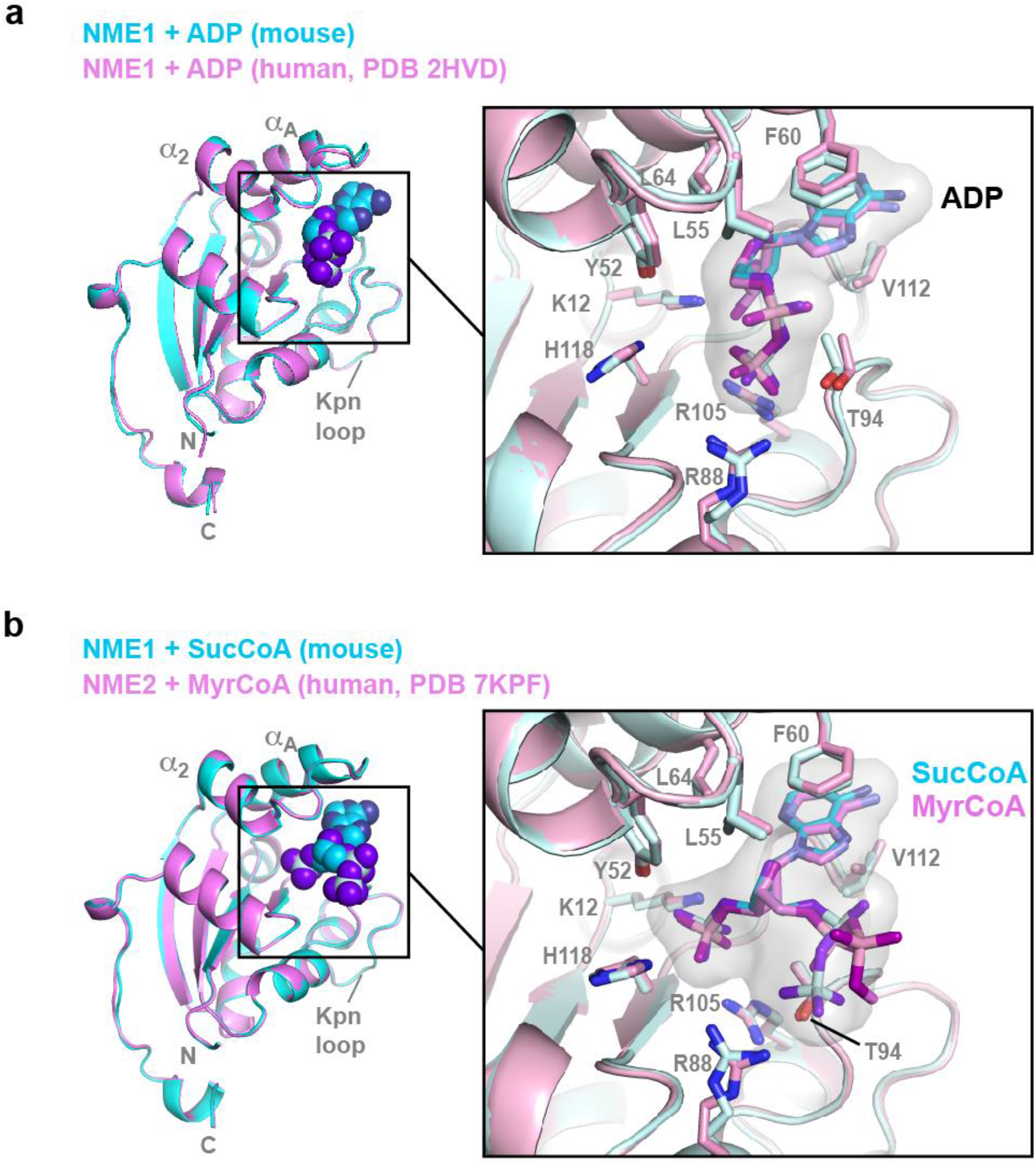
Similarity with previously reported NME1/2 structures. **a.** Alignment of ADP-bound structures of murine NME1 and human NME1 (PDB 2HVD)^26^. **b.** Structural alignment of murine NME1 bound to SucCoA with human NME2 bound to myristoyl-CoA (PDB 7KPF)^15^.

**Figure S5.**
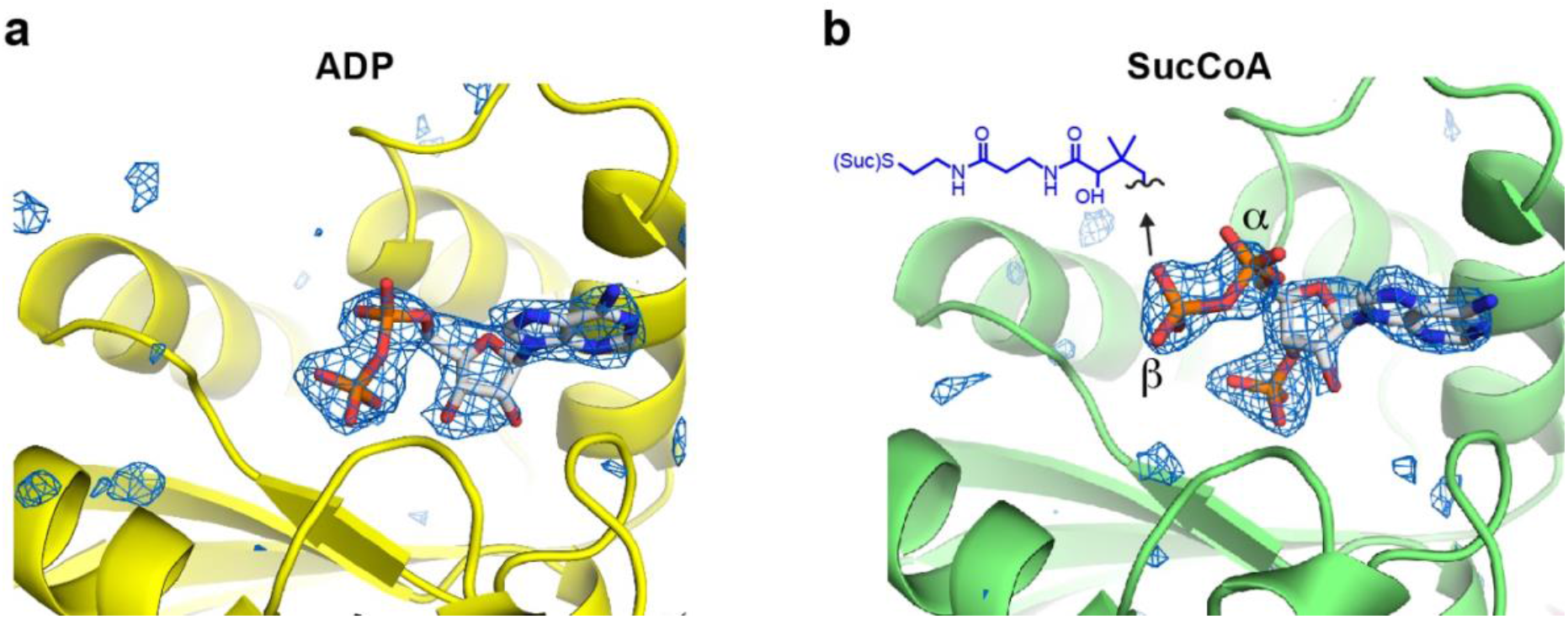
Omit map density for (A) ADP and (B) SucCoA bound to NME1. The electron density (mesh in blue) shows *F_o_*-*F_c_* omit maps contoured at 3.0σ (ADP) and 2.0σ (SucCoA) where ligands were omitted from the map calculation. No strong density is observed for the pantetheine and succinyl moities of the SucCoA ligand.

**Figure S6.**
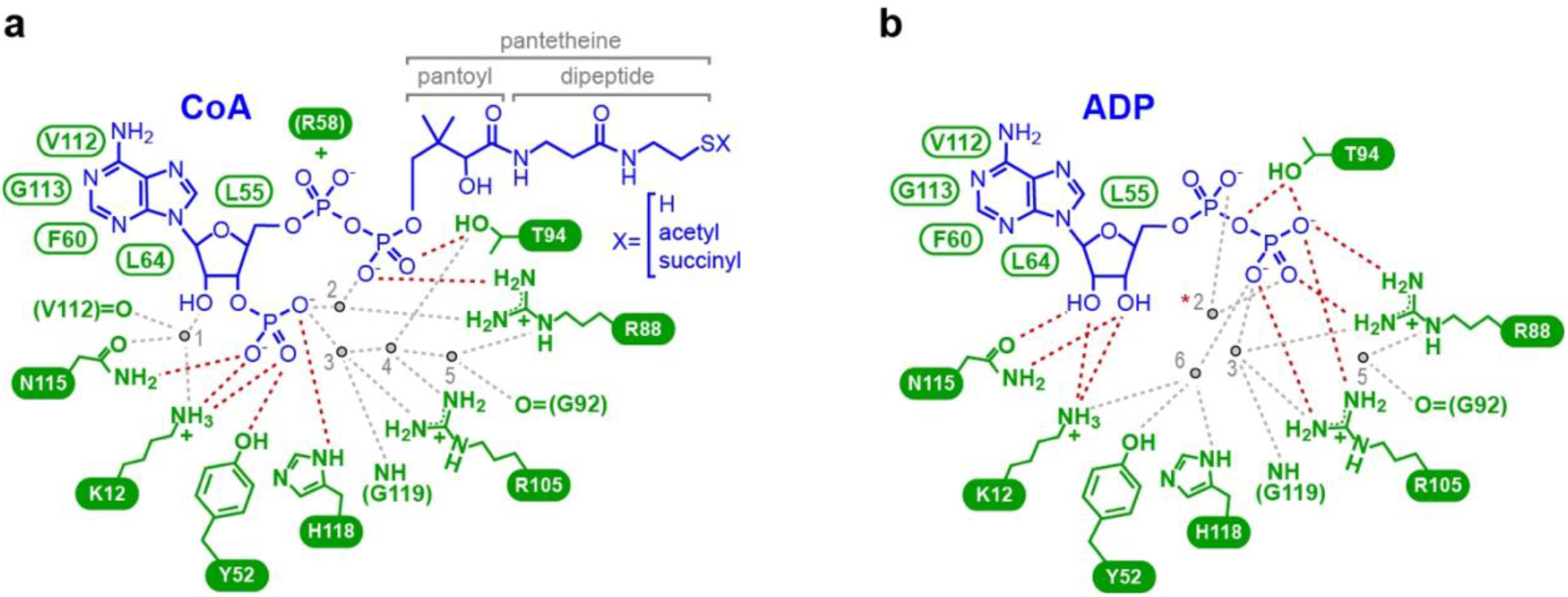
Schematic summary of NME1 interactions mediating recognition of (a) CoA/SucCoA and (b) ADP. Direct and water-mediated hydrogen bonds are shown as red and gray dashed lines, respectively. NME1 residues that mediate H-bonding are indicated in white on a green background; residues mediating van der Waals contacts or aromatic stacking interactions are labelled in green. The Arg58 residue which forms a salt bridge with the CoA α-phosphate in a subset of NME1 subunits is shown in parentheses. Structurally conserved water molecules are numbered 1-6. Three of these are conserved between the ADP- and SucCoA-bound structures. In some previously reported ADP-bound NDPK structures (PDB 1NUE and 1NDP) a Mg^2+^ ion replaces water molecule 2 (red asterisk). The CoA 3’ phosphate group forms direct H bonds with Lys12, Tyr52, Asn115 and His118 as well as water-mediated H bonds with Arg88, Arg105 and the Gly119 backbone. In the ADP-bound structure these residues either hydrogen bond with the ribose 2’ and 3’ OH groups or with a water molecule H-bonded to the β-phosphate. Because the CoA ribose is displaced away from the protein by the 3’ phosphate, the H bonds that NME1 residues Lys12 and Asn115 make with the sugar 2’ OH group of ADP cannot be maintained. These are instead replaced by water-mediated bonds that allow the larger protein-ligand distance to be bridged. Similarly, the outward shift of the CoA β-phosphate is accommodated by water molecules that satisfy the H bonding potential of residues Arg88 and Arg105, which in the ADP-bound structure directly contact the β-phosphate.

**Figure S7.**
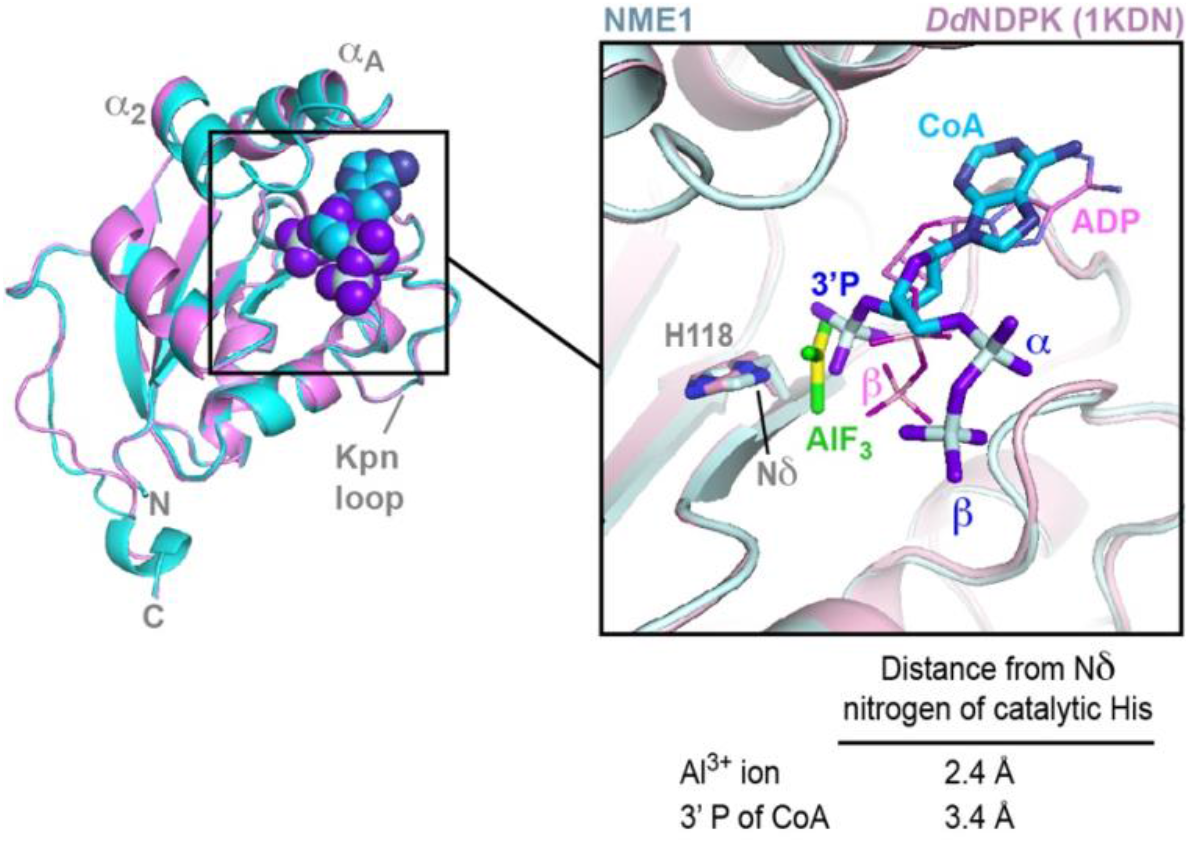
The CoA β-phosphate occupies the position of the ATP γ-phosphate prior to nucleophilic attack. Alignment of the CoA-bound NME1 structure with that of *Dd*NDPK bound to ADP-AlF_3_, solved at 2.8 Å-resolution^67^. AlF_3_ mimics the transition state of the ATP γ phosphate. The ADP ligand bound to *Dd*NDPK is shown with a thin stick radius. The CoA 3’ phosphate is within 1.5 Å of the aluminum ion in AlF_3_. Compared to the Al^3+^ ion, the CoA 3’ phosphate is farther from the histidine’s nucleophilic Nδ nitrogen and closer to the β-phosphate of ADP-AlF_3_, suggesting that the CoA 3’ phosphate occupies the position of the ATP γ-phosphate prior to nucleophilic attack.

**Figure S8.**
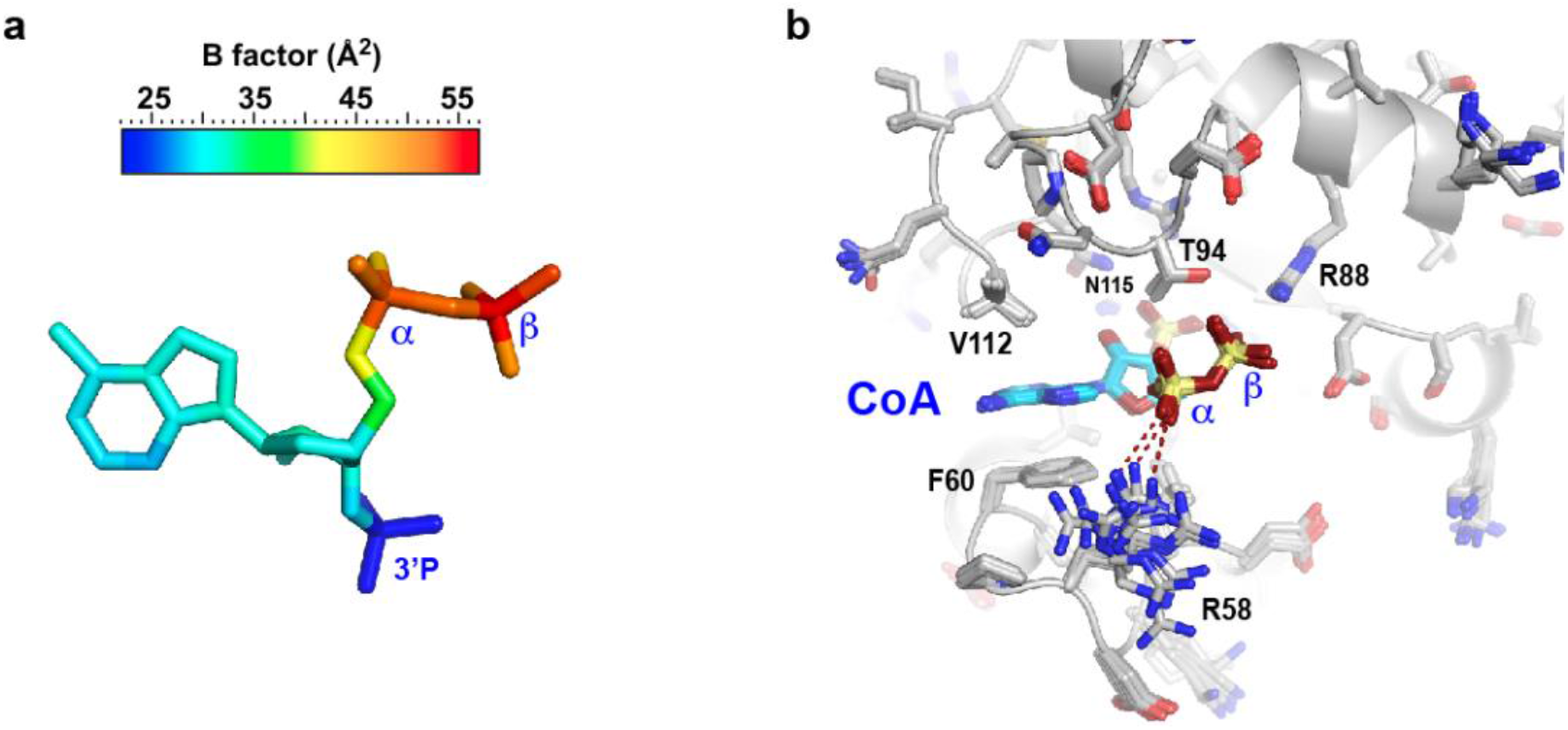
The α- and β-phosphates of the CoA nucleotide are poorly recognized by NME1. **a.** Structure of the NME1-bound CoA nucleotide with atoms colored according to their average crystallographic B factor, showing that the α- and β-phosphates are the most mobile atoms in the ligand. In contrast, the 3’ phosphate is highly constrained by numerous H bond interactions with the protein, explaining the low values of its B factors. The average B factors shown were calculated by averaging over the 12 independent copies of the ligand in the crystal structure of SucCoA-bound NME1. **b.** Alignment of the 12 crystallographically independent subunits of NME1 in the crystal structure of SucCoA-bound NME1. Active site residues adopt a uniform sidechain conformation in all 12 subunits except for residue Arg58, which is located close to the CoA α-phosphate and exhibits considerable variability. Arg58 is within H bonding distance of the CoA α-phosphate in only 4 of the 12 monomers. H bonds are indicated as red dashed lines.

**Figure S9.**
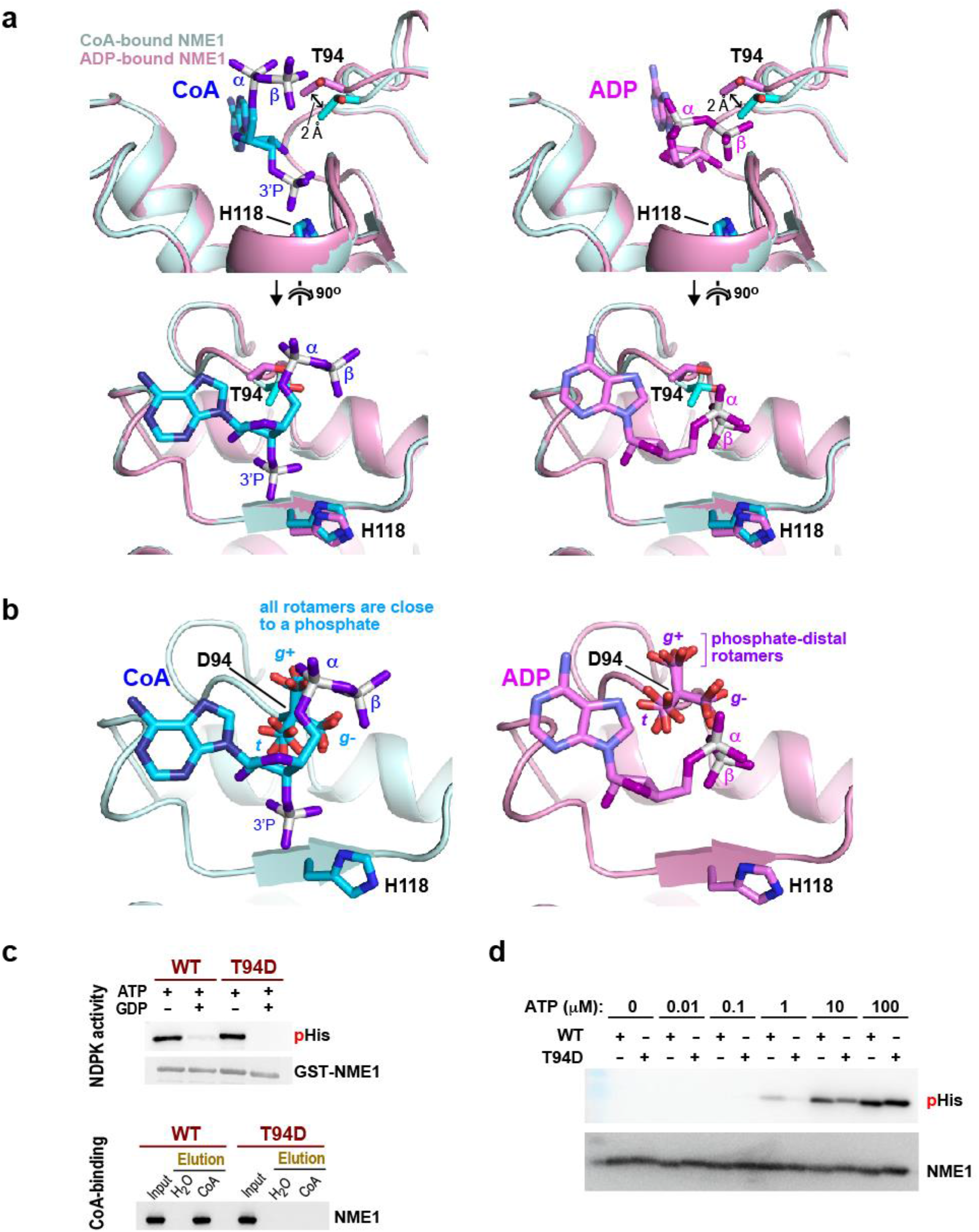
Positional shift of Thr94 and analysis of the T94D mutant. **a.** Structural alignment of the CoA- and ADP-bound NME1 illustrating the shift of position of residue Thr94. Only one ligand, CoA or ADP, is shown in the left and right views, respectively. Thr94 is the only active site residue whose backbone atoms shift significantly between the CoA- and ADP-bound structures, resulting in a ∼2 Å displacement of the sidechain hydroxyl and methyl groups. The lower view shows that these groups sit directly below the diphosphate of CoA but are more peripherally located with respect to that of ADP. **b.** *In silico* modeling of the T94D mutant. The Asp residue is modeled in its nine preferred sidechain conformations. The labels *g^+^, g^-^* and *t* refer to the *gauche^+^, gauche^-^* and *trans* conformations of the χ_1_ angle. *Left:* Replacing the Thr residue in the CoA-bound NME1 crystal structure by an Asp residue and modeling its sidechain conformations revealed that all Asp rotamers point their negatively charged carboxylate group towards one or more CoA phosphate groups located within unfavourably close proximity. *Right:* Repeating the same exercise with the ADP-bound structure revealed several Asp rotamers (all with χ_1_ in a *g^+^* conformation) that point their carboxylate away from the ADP diphosphate, located over 5 Å away. This suggests that replacing Thr94 by an Asp residue should yield an NME1 mutant that could accommodate ADP in the active site by adopting a permissive Asp94 rotamer, whereas no Asp94 conformation could avoid an electrostatic repulsion with a bound CoA ligand. **c.** *Upper panel.* The NME1 T94D mutant retains significant NDPK activity. Both WT and mutant forms of GST-tagged NME1 are phosphorylated on histidine after incubation with ATP (100 μM) and become dephosphorylated upon incubation with GDP (200 μM). Proteins were detected using anti-phosphohistidine (pHis) and anti-NME1 antibodies (NME1). *Lower panel.* The T94D mutant is defective for CoA-binding activity. WT and mutant NME1 were incubated with CoA beads, eluted with free CoA after a pull-down as described in **Fig. 1b** and the blot was probed with an anti-NME1 antibody. **d.** The T94D mutant has lower NDPK activity than WT. Both WT and mutant forms of NME1 are phosphorylated on histidine following incubation with ATP. The degree of phosphorylation observed at 1-10 μM ATP is approximately an order of magnitude lower for the mutant compared to the WT. Proteins were detected using anti-phosphohistidine (pHis) and anti-NME1 antibodies (NME1).

**Figure S10.**
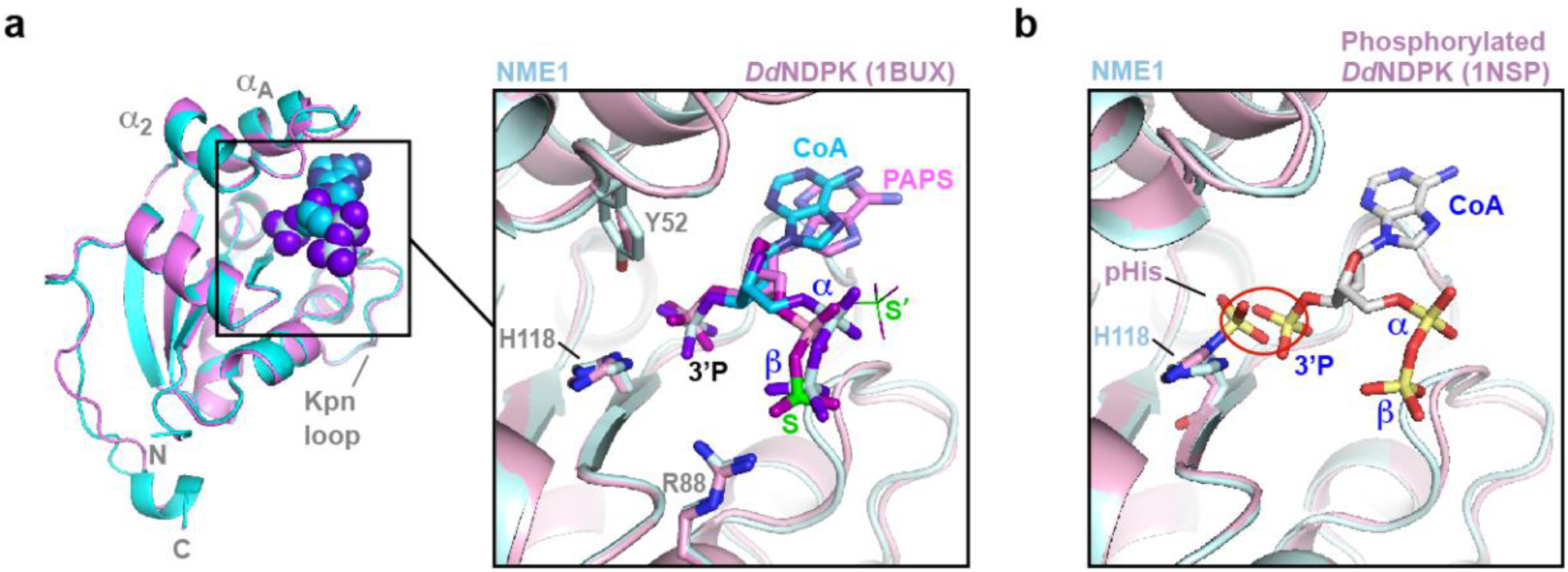
Structural analyses predict that CoA binding and histidine phosphorylation are mutually inhibitory. **a.** Alignment of SucCoA-bound NME1 and PAPS-bound *Dd*NDPK (PDB 1BUX). PAPS is chemically identical to the CoA nucleotide except that its sulfate group replaces the CoA β-phosphate. In the crystal structure, the PAPS sulfate group was observed to occupy two positions, each with partial occupancy, labeled S and S’. The latter is shown with a thinner stick radius. Residue numbering is that of mammalian NME1. **b.** Structural alignment of NME1:SucCoA with *Dd*NDPK (the drosophila NDPK) phosphorylated on His119 (equivalent to murine His118) (PDB 1NSP) predicts a steric clash and electrostatic repulsion between the histidine phosphate and the CoA 3’ phosphate, located only 1.4 Å apart (red oval).

**Figure S11.**
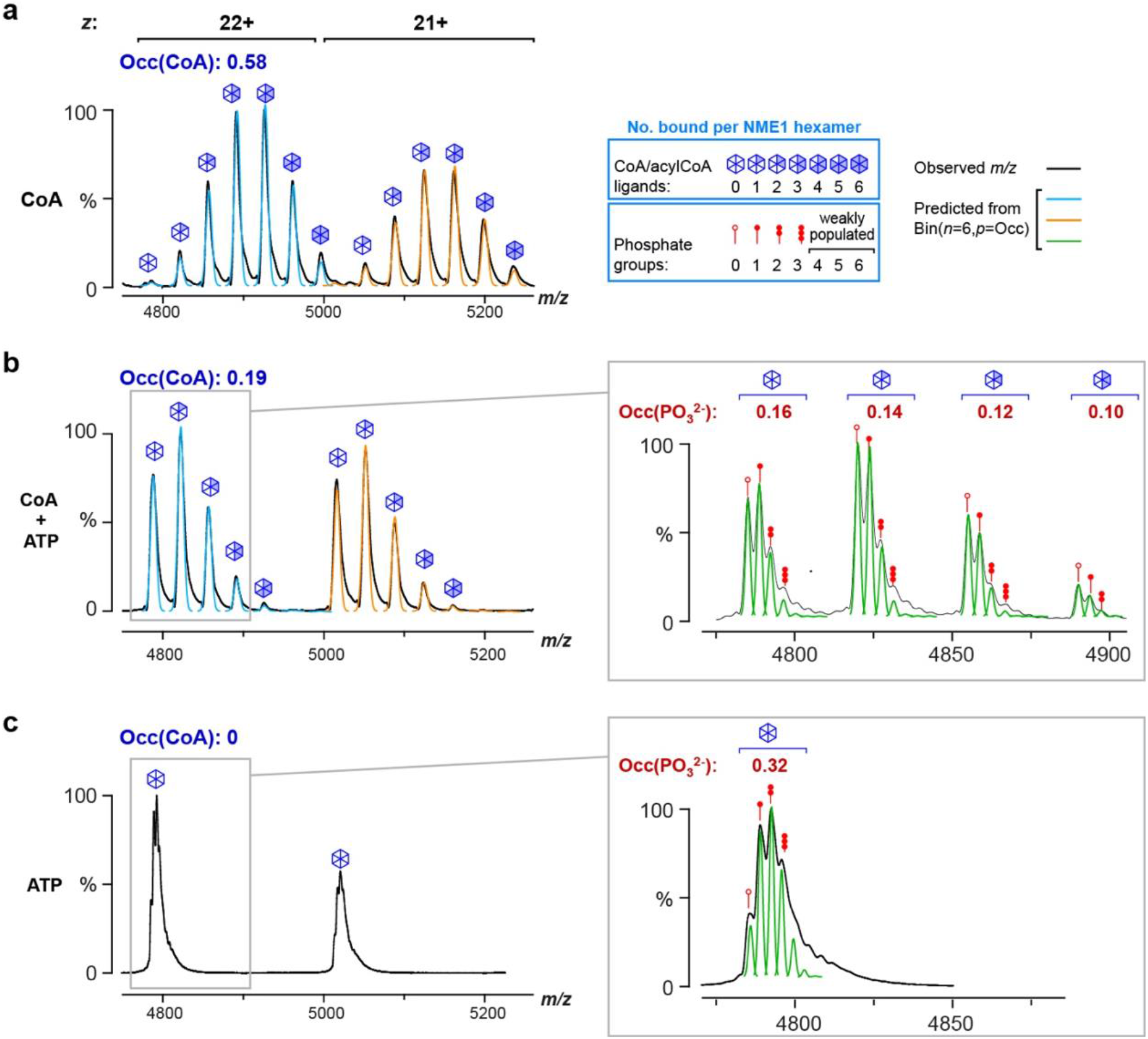
CoA binding and NME1 phosphorylation are mutually inhibitory. Native MS spectra shown are the same as the corresponding spectra in **Fig. 2e**. Primary peaks in the spectra were modelled as a Gaussian function convoluted with a binomial distribution (blue and orange curves) to describe the distribution of CoA-bound NME1 states, as described in **Fig. S3**. *Insets*. A magnified view of individual peaks reveals sub-peaks corresponding to the presence of phosphoryl groups on NME1. The distribution of phosphorylation states was fitted as a Gaussian function convoluted with a binomial distribution (green curves) in a manner analogous to that used for primary peak fitting. The reasonable agreement between the recorded spectra and the fitted curves confirms that phosphorylation occurs independently at the six catalytic sites of the NME1 hexamer, allowing one to estimate the fraction of NME1 monomers that are phosphorylated [*Occ*(PO_3_^2-^)].

**Figure S12.**
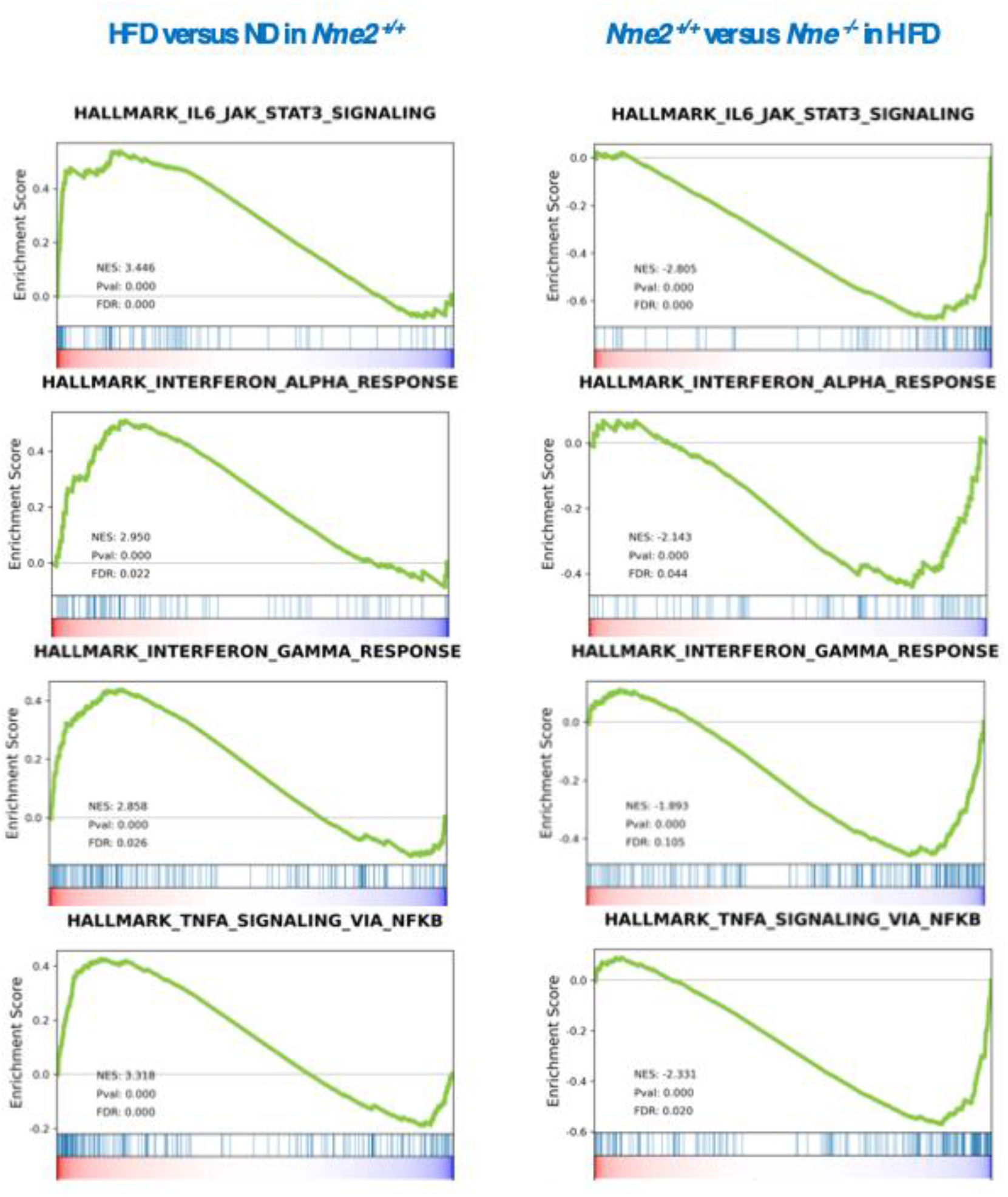
NME2-dependent activation of genes involved in protective signaling pathways in the liver of HFD mice. GSEA plots showing that gene sets corresponding to genes involved in the indicated signaling pathways are enriched in the hepatocytes of WT mice treated with HFD for 6 weeks compared to ND (left panels). These same gene sets are depleted in the liver of *Nme2^-/-^* mice under HFD compared to *Nme2^+/+^* mice, demonstrating that, in the absence of NME2, the corresponding signaling pathways are not activated under HFD.

**Figure S13.**
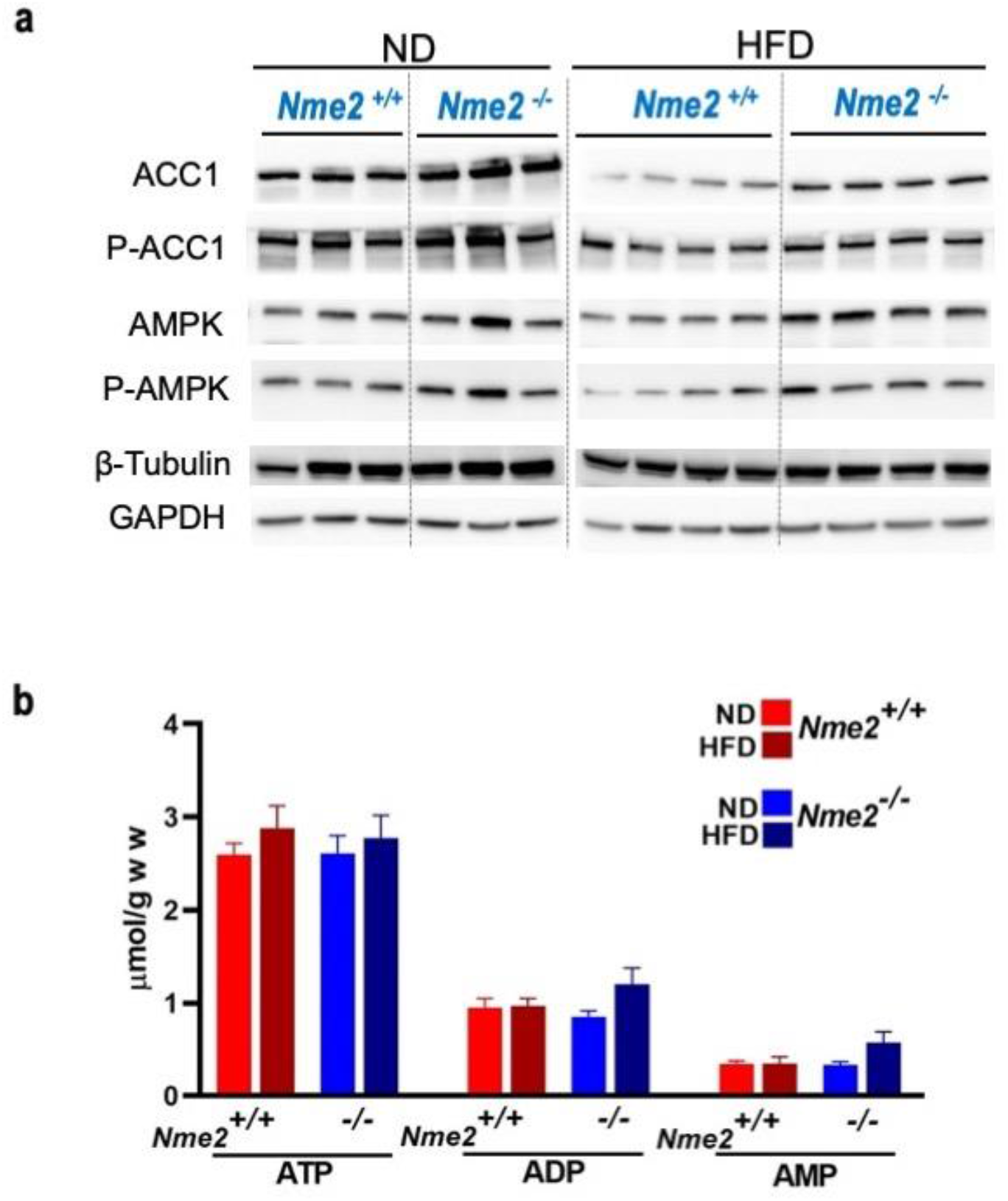
AMPK signaling is not involved in the response to a HFD challenge and the NME2- dependent control of fatty-acid synthesis. **a.** Extracts from livers from three independent *Nme2^+/+^* or *Nme2^-/-^* mice under ND or from four independent *Nme2^+/+^* or *Nme2^-/-^* mice under HFD for 6 weeks were probed with the indicated antibodies. **b.** Cellular concentrations of ATP, ADP and AMP measured in liver extracts from *Nme2^+/+^* and *Nme2^-/-^* mice. For each measurement, livers from independent mice were used as follows: *Nme2^+/+^* ND, n = 9; *Nme2^+/+^* HFD, n = 4; *Nme2^-/-^* ND, n = 9; *Nme2^-/-^* HFD, n = 5. The graph shows the average values (in μmol/g of liver) and SEM.

## RESOURCES TABLE

### Key resources table

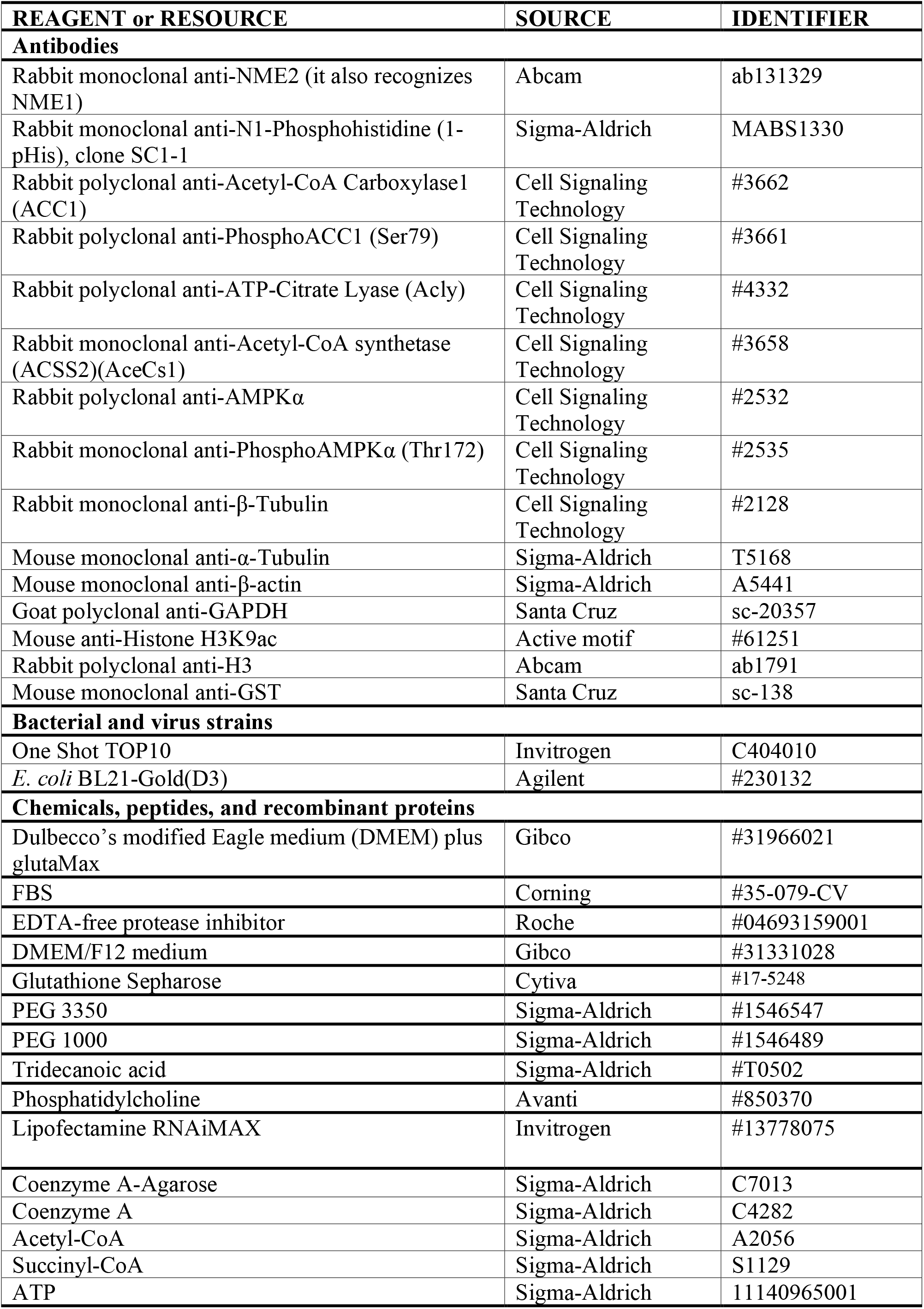
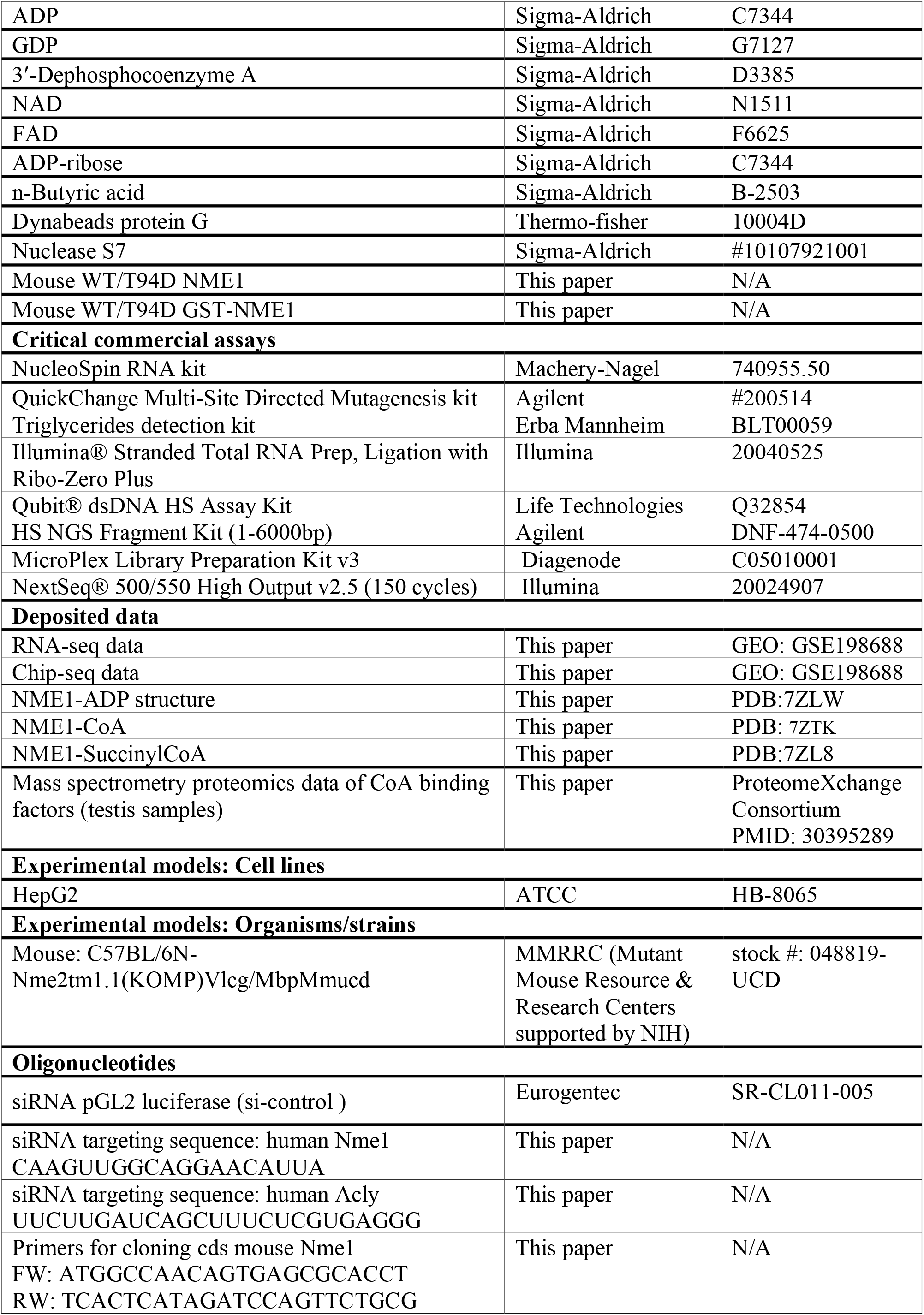
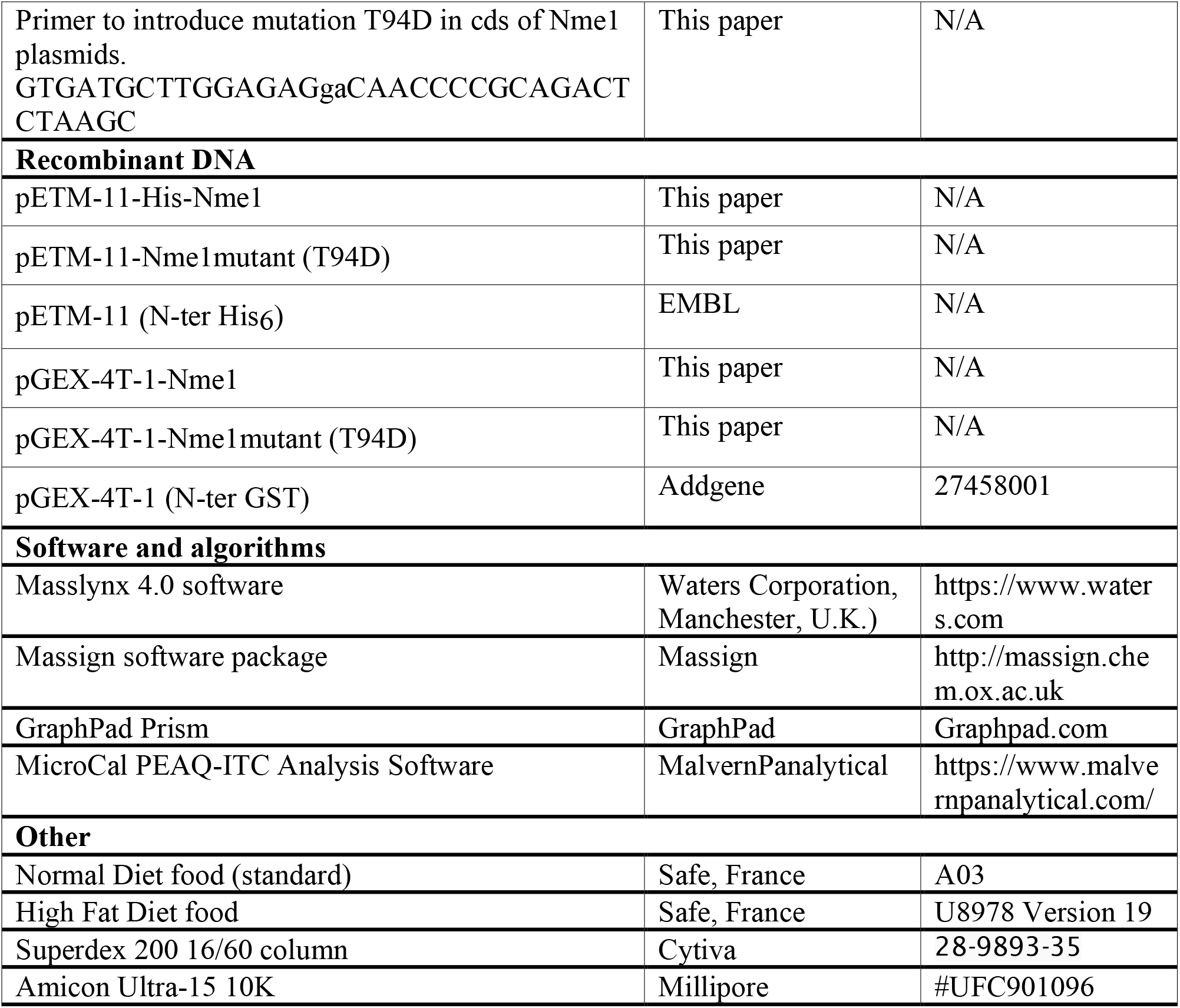

## RESOURCE AVAILABILITY

### Lead contact

Further information and requests for resources and reagents should be directed to and will be fulfilled by the lead contact, Saadi Khochbin.

### Material availability

Plasmids generated and detailed in this manuscript are freely available for academic use from the Khochbin Lab.

### Data and code availability

The mass spectrometry proteomics data of CoA binding factors (testis samples) have been deposited to the ProteomeXchange Consortium via the PRIDE^68^ partner repository with the dataset identifier PXD029159 and 10.6019/PXD029159 (https://www.ebi.ac.uk/pride/login (Username; Password: 3JlxQLlB)).

Crystal structures and diffraction data for the NME1 protein bound to ADP (PDB 7ZLW), CoA (PDB: 7ZTK) and SucCoA (PDB 7ZL8) have been deposited in the Protein Data Bank (www.rcsb.org). RNA-seq and ChIP-seq data are deposited in GEO SuperSeries accession # GSE198688.

To review GEO accession GSE198688:

### Go to https://www.ncbi.nlm.nih.gov/geo/query/acc.cgi?acc=GSE198688 Enter token wtajyaiwbjgdzmr into the box

All other data are available upon request to the corresponding authors.

## EXPERIMENTAL MODEL AND SUBJECT DETAILS

### Escherichia coli culture

*E. coli* BL21-Gold(D3) (Agilent #230132) bacteria were transformed by pETM-11-Nme1/T94D- mutant and pGEX-4T-1-Nme1/T94D-mutant and clones were selected on LB agar plates containing kanamycin and ampicillin, respectively, and cultured at 37°C.

One Shot TOP10 Chemically Competent *E. coli* (Invitrogen #C404010) were cultured at 37°C to amplify the plasmids pETM-11-Nme1/T94D-mutant, and pGEX-4T-1-Nme1/T94D-mutant.

### Animals

All mouse experiment protocols were approved by the official ethics committee of the University Grenoble Alpes (ComEth, C2EA-12) and all the investigators directly involved in the care and breeding of the mice have an official animal-handling authorization obtained after 2 weeks of intensive training and a final formal evaluation. All mice were in a light-regulated colony room (12 hours light/12 hours dark), with food and water available ad libidum. Heterozygotes Nme2 KO male and female mice (C57BL/6N-Nme2tm1.1(KOMP)Vlcg/MbpMmucd) were obtained from MMRRC (Mutant Mouse Resource & Research Centers supported by NIH) (stock #: 048819-UCD). Mice were bred in animal core facility (Grenoble High Technology Animal Facility (PHTA), University Grenoble Alpes). Both male and female mice of 2 to 5 months of age were used for all studies.

### Mammalian Cells

HepG2 cells (ATCC #HB-8065) were cultured at 37°C in Dulbecco’s modified Eagle medium (DMEM) plus glutaMax^TM^ (Gibco #31966021) supplemented with 10% fetal bovine serum (FBS) (Corning #35-079-CV).

## METHOD DETAILS

### Lysate preparation for CoA pulldown Total testis

For each CoA pull-down experiment, two adult male mice were euthanized and the testes collected. After removing the albuginea, the seminiferous tubules were directly immersed in 1 ml of lysis buffer (20% glycerol, 3 mM MgCl_2_, 50 mM Hepes, 250/500 mM KCl, 0.1% NP40, 1 mM DTT, 1 mini-tablet of EDTA-free protease inhibitor (Roche #04693159001)), homogenized for up to one minute with homogenizer (Heidolph RGL 55/1) and incubated for 30 min on ice. The sample was then centrifugated at 394 g for 10 min at 4°C and the supernatant extract was used immediately for the purification of the CoA bound proteins by CoA pull-down. The 500 mM KCl lysate was diluted to 250 mM with buffer before proceeding.

### Spermatogenic cells

The enrichment (or fractionation) of spermatogenic cells at different stages of their maturation from the testes of 6 mice, was performed as described in detail in refs.^69, 70^. Briefly, the seminiferous tubules were incubated with 1 mg/ml collagenase/PBS for 15 min at 35°C, dissociated by pipetting for 10 min and filtered through a 100 µm filter. Total germ cells were pelleted, suspended in 0.5% BSA-DMEM/F12 medium (Gibco #31331028) and sedimented through 2%-4% BSA gradient for 70 min at 4°C. The fractions were than collected and examined under a phase contrast microscope. Fractions of Total germ cells (TGC, spermatogenic cells before sedimentation), or respectively enriched in Pachytene Spermatocytes (PS), in Round Spermatids (RS) as well as in Elongating and Condensing Spermatids (ECS) were pooled for the preparation of cell lysates. In each CoA pull-down experiment, the cell pellets were incubated in 125 µl of lysis buffer (20% glycerol, 3 mM MgCl_2_, 50 mM Hepes, 500 mM KCl, 0.1% NP40, 1 mM DTT, 1 mini-tablet of EDTA-free protease inhibitor) and treated as previously described for total testis lysate. **Liver** 50 mg of liver from *Nme2* WT or *Nme2* KO mice were incubated in 500 μl of 250 mM lysis buffer and treated as previously described for total testis lysate.

### Purification of CoA-bound proteins

Coenzyme A-Agarose (C7013-500MG, Sigma-Aldrich) beads were suspended in 3.5 ml of 10 mM sodium acetate (pH 6.0) and stored at -20°C. For each experiment, 50 µl of the CoA-agarose mixture was washed 3 times in lysis buffer (250 mM KCl) and mixed with 1 ml of each of the lysates (respectively from testis, spermatogenic cells or liver) and incubated with rotation overnight at 4°C. The CoA-agarose beads were then washed three times in the lysis buffer, followed by a final wash with PBS. Half of the beads were then incubated for 1 h at 4°C in 50 µl of 5 mM free CoA (C4282, Sigma-Aldrich) (dissolved in H_2_O), and the other half was incubated in 50 µl of H_2_O (Control). Eluted proteins were collected following a centrifugation step (1650 g, 2 min, 4°C). 10 µl of the eluted proteins were separated by migration on a NuPAGE 4–12% Bis-Tris Protein Gel (Invitrogen) and silver stained. Another 10 µl were analyzed by Western Blotting. The rest of the eluted proteins were analyzed by Mass Spectrometry (MS).

### Mass spectrometry (MS)-based proteomic analyses of the CoA bound proteins

The eluted proteins solubilized in Laemmli buffer were stacked in the top of a 4-12% NuPAGE gel (Invitrogen). After staining with R-250 Coomassie Blue (Biorad), the proteins were digested in-gel using trypsin (modified, sequencing purity, Promega), as previously described^71^. The resulting peptides were analyzed by online nanoliquid chromatography coupled to MS/MS (Ultimate 3000 and LTQ-Orbitrap Velos Pro, Thermo Fisher Scientific) using a 124 min gradient. For this purpose, the peptides were sampled on a precolumn (300 μm x 5 mm PepMap C18, Thermo Scientific) and separated in a 75 μm x 250 mm C18 column (PepMap C18, 3 μm, Thermo Fisher Scientific). The MS and MS/MS data were acquired by Xcalibur (Thermo Fisher Scientific).

The peptides and proteins were identified by Mascot (version 2.7.0, Matrix Science) through concomitant searches against the Uniprot database (*Mus musculus* taxonomy, June 2021 download), and a homemade classical database containing the sequences of classical contaminant proteins found in proteomic analyses (human keratins, trypsin, etc.). Trypsin/P was chosen as the enzyme and two missed cleavages were allowed. Precursor and fragment mass error tolerances were set respectively at 10 ppm and 0.6 Da. Peptide modifications allowed during the search were: Carbamidomethyl (C, fixed), Acetyl (Protein N-term, variable) and Oxidation (M, variable). The Proline software^72^ was used for the compilation, grouping, and filtering of the results (conservation of rank 1 peptides, peptide length ≥ 6 amino acids, false discovery rate of peptide-spectrum-match identifications < 1%, and minimum of one specific peptide per identified protein group^73^. Proline was then used to perform MS1-based label free quantification of the identified protein groups based on razor and specific peptides. The proteins from the contaminant database were discarded from the final list of identified proteins. To calculate fold changes between CoA and control eluates, missing values were imputed in ProStaR ^74^ using the DetQuantile algorithm (imputation with a value corresponding to the first percentile of the sample). To be considered as a potential CoA interactor, a protein must be quantified with a minimum of two peptides and be enriched at least 10 times in the CoA eluates compared to the control eluate. The relative abundance of the different proteins in each sample was evaluated through calculation of their intensity-based absolute quantification (iBAQ)^54^ values.

### Gene cloning

The full-length codon sequence of mouse *Nme1* (CCDS 25247.1) was cloned in pGEX-4T-1 (GST tag) and pETM-11 (His tag) vectors for protein production in BL21 *E. coli*. The T94D mutation was introduced in the plasmids by PCR using the QuickChange Multi-Site Directed Mutagenesis kit (#200514 -Agilent Technologies). The sequence of the primer used to introduce the mutation T94D in the *Nme1* sequence of plasmids is reported in key resources table. Every plasmid was sequence-verified.

### Recombinant mouse GST-NME1 purification

*Escherichia coli* BL21-Gold(D3) (230132, Agilent) were transformed with 50 ng of either GST- Nme1 or GST- NME1-T94D-mutant vector (pGEX-4T-1) and cultured from a single colony. Once 600 ml of culture reached an optical density at 600 nm (OD_600nm_) of 0.8, protein expression was induced by adding 0.2 mM IPTG, followed by incubation with shaking overnight at 18°C. The cells were pelleted at 9000 g at 4°C for 30 min, resuspended in 10 ml of 50 mM Tris-HCl pH 7.4, 500 mM NaCl, 0.3 % Triton plus protease inhibitors, and disrupted by sonication for 2 min using cycles of alternating on/off pulses (5 s On/5 s Off) at 4°C. Then, the lysate was clarified by centrifugation at 48,000 g 4°C for 20 min, and incubated for 1 h at room temperature with 2 ml of pre-washed Glutathione Sepharose (Cytiva Cat#17-5132-01). After three washes with a buffer containing 50 mM Tris-HCl pH 7.4, 500 mM NaCl and protease inhibitors, GST-NME1 was eluted three times for 30 min each in 1 ml of a buffer containing 50 mM Tris-HCl pH 7.4, 500 mM NaCl and 25 mM glutatione. The GST-NME1 proteins (WT and T94D-mutant) were collected and adjusted to 1 mg/ml in the same buffer, and subsequently flash-cooled in liquid nitrogen for storage at -80°C.

### Recombinant mouse NME1 purification for biochemical and structural analyses

*Escherichia coli* BL21-Gold(D3) (230132, Agilent) were transformed with 50 ng of the His- Nme1/T94D-mutant vector (pETM-11) and cultured from a single colony. Once 1 L of culture reached an OD_600nm_ of 0.8, protein expression was induced with 0.2 mM IPTG and the culture was incubated with shaking overnight at 18°C. The cells were pelleted by centrifugation at 9000 g at 4°C for 30 min and stored at -80°C until further use. The pellet was subsequently thawed, resuspended in 12.5 ml of phosphate buffer 0.1 M, pH 7.4, containing 150 mM NaCl, 25 mM imidazole and protease inhibitors (buffer A) and the cells disrupted by sonication, using 60 cycles of alternating on/off pulses (2 s on/10 s off) at 4°C. Then, the lysate was clarified by centrifugation at 48,000 g at 4°C for 1 h, and applied to a 5 ml HisTrap HP prepacked column (Cytiva #175248). The column was then washed with buffer A (15 to 20 column volumes), NME1 was eluted in a 100 ml linear gradient ranging from 0 to 500 mM imidazole using 0.1 M phosphate buffer pH 7.4, 150 mM NaCl, 500 mM imidazole (buffer B) and collected. The N-terminal 6-His tag was cleaved by TEV protease (1 mg protease per 15 mg His-NME1) during dialysis against phosphate buffer with 150 mM NaCl overnight at 4°C. After removal of the cleaved His-tag by a 5 ml HisTrap HP column, NME1 was collected from the flow-through and subjected to size exclusion chromatography using a Superdex 200 16/60 column (Cytiva) which was previously equilibrated with phosphate buffer containing 150 mM NaCl. NME1 appeared at an elution volume of approximately 170 ml. The protein was collected, pooled and concentrated to 11 mg/ml using a 10-kDa cut-off centrifugal filter (Amicon® Ultra-15 10K, Millipore #UFC901096), flash-cooled in liquid nitrogen and stored at -80°C.

### Crystallization and crystal structure determination

Protein crystallization was performed by the hanging drop vapor diffusion method at 20°C (for ADP- and CoA-bound NME1) or 4°C (for SucCoA-bound NME1) by mixing 1 μl of the NME1/ligand complex with 1 μl of reservoir solution. ADP-bound NME1 was crystallized by mixing a solution of 6.1 mg/ml NME1, 1 mM ADP, 250 mM NaCl and 25 mM Tris pH 7.4 with 0.2 M NaCl, 0.1 M Tris pH 8.0 and 30% (w/v) PEG 400. CoA-bound NME1 was crystallized by mixing a solution of 5.5 mg/ml NME1, 1 mM CoA, 150 mM NaCl and 0.1 M phosphate buffer pH 7.4 with 0.2 M Mg(NO_3_)_2_ and 36% PEG 3350. SucCoA-bound NME1 was crystallized by mixing a solution of 5.5 mg/ml NME1, 1 mM SucCoA, 150 mM NaCl and 0.1 M phosphate buffer pH 7.4 with 0.1 M citric acid pH 3.5 and 4-10% PEG 1000. Harvested crystals were flash-cooled in liquid nitrogen.

Diffraction data were collected at beamline ID30A-1 of the European Synchrotron Radiation Facility (ESRF). All crystals were monoclinic with space group P2_1_ and contained either one (ADP, CoA) or two (SucCoA) NME1 hexamers in the asymmetric unit. Data collection statistics are summarized in **Table S2**. Data for ADP- and SucCoA-bound NME1 were automatically integrated with XDS^75^ which is included in the automatic software toolbox auto-PROC^76^ and scaled with AIMLESS^77^. Data for CoA-bound NME1 was manually integrated with XDS and scaled with AIMLESS.

Structures were solved by molecular replacement in Phaser^78^ using the structure of human NME1 bound to ADP (PDB 2HVD)^26^ or imidazole fluorosulfate (PDB 5UI4)^79^ after removing ligands and water molecules, as search models. Iterative rounds of refinement and model building were performed using PHENIX^13^ and Coot^80^, resulting in the final R-values shown in **Table S2**. In the ADP-bound structure, ADP is observed bound to 5 of the 6 subunits in the NME1 hexamer, whereas the remaining binding site is empty because of crystal packing constraints. For CoA- and SucCoA-bound NME1 all ligand-binding sites are occupied. Weak or missing electron density precluded reliable modeling of the CoA/SucCoA pantetheine moiety and the SucCoA succinyl group, which are consequently omitted from the final refined atomic coordinates.

### *In vitro* histidine phosphorylation assay

*In vitro* autophosphorylation of the WT or T94D mutant forms of untagged or GST-tagged NME1 was performed in TMD buffer (20 mM TRIS-HCl pH 8.8, 5mM MgCl_2_, and 1 mM DTT) with ATP at room temperature in a 20 µl reaction volume. In reaction reported in **Fig. S2b**, 0.23 mM of NME1 was incubated with 0.5 mM ATP for 10 min and subsequently 0.028 mM of phosphorylated NME1 was incubated with GDP (0.1, 0.2 and 0.4 mM) for other 10 min. In **Fig. S2a**, 5.7 mM of NME1 was incubated with 100 μM ATP for 10 min. In **Fig. 2c**, GST-WT and T94D mutant (1 μg) were incubated with 100 μM ATP for 10 min and subsequently with GDP (200 μM) for other 10 min. In **Fig. S9c**, WT and T94D mutant (1 μg) were incubated with ATP (0-100 μM) for 1 min. Each reaction was arrested by adding 5 μL of 5X gel loading buffer (10% SDS, 250 mM Tris-HCl pH 8.8, 0.02% bromophenol blue, 50% glycerol, 50 mM EDTA, 500 mM DTT) and 10 μl of the solution was analyzed immediately by western blotting to detect 1-*P*-histidine phosphorylation. The western blot was performed as reported in detail in ref.^81^ with some modifications. Briefly, the stacking gel buffer was adjusted to pH 8-9 and the samples were not heated. All electrophoresis steps were performed at 4°C and samples were resolved at 100 V for 2-3 h. Proteins were transferred to PVDF membranes at 100 V for 2 h at 4°C and immediately incubated for 1 h at room temperature in BSA Blocking Buffer 3% BSA in 1X TBS-T (20 mM Tris buffer pH8.8, 150 mM NaCl, 0.1% tween 20). The membranes were incubated with a primary Anti 1-*P*-Histidine antibody (clone SC1-1 1/1000 in TBS-T) were incubated on the membrane overnight at 4°C. After incubation with HRP-conjugated secondary antibodies, the membranes were washed three times for 10 min each with 0.1% TBS-T. The revelation of the membranes was performed using the Clarity Western ECL Blotting Substrate (Biorad 1705060) following the usual procedure.

### Native PAGE

NME1 (24 μM) was incubated for 10 min at RT with ATP (from 0 to 384 μM) and phosphorylated NME1 (previously prepared by incubating with 192 μM ATP) was incubated with ADP (from 192 to 1728 μM) in TMD buffer for 10 min at RT. After adding 2 μL of native loading buffer (62.8 mM Tris HCl pH 6.8, 40 % glycerol, 0.01 % bromophenol blue), 5 μL of samples were analysed on a 10% TGX (Biorad) gel, run under native conditions (0.5X TBE buffer, 4°C, 100 V, 120 min). Gels were stained with Coomassie blue and scanned on a ChemiDoc MP gel imaging system (Biorad).

### Nano-differential scanning fluorimetry (nano-DSF)

10 μL samples of 0.5 mM NME1 prepared in the absence or presence of either CoA, AcCoA, SucCoA or ADP at a concentration of 0.62 mM in a buffer containing 25 mM Tris pH 7.4, 250 mM NaCl were loaded into nanoDSF Grade Standard Capillaries (NanoTemper, #PR-C002) and analysed on a Prometheus NT 48 instrument (Nanotemper). Assay samples were heated from 20°C to 95°C at a rate of 0.5°C/min. Intrinsic tryptophan fluorescence was measured at 330 nm and 350 nm using an excitation power of 10%. The 350/330 fluorescence ratio was plotted against temperature and the T_m_ value determined from the inflection point using the instrument software.

### Isothermal titration calorimetry (ITC)

Calorimetric experiments were performed in triplicate on a MicroCal iTC200 calorimeter (Malvern Panalytical) at 20°C while stirring at 330 rpm. The syringe and cell were respectively filled with 900 µM CoA and 30 µM NME1, both in 0.1 M phosphate buffer pH 7.4, 150 mM NaCl. Titrations consisted of 50 identical injections of 6 µl made at time intervals of 450 s. ITC data were corrected for the heating of CoA injection into buffer and analyzed with the MicroCal PEAQ-ITC Analysis Software (Malvern Panalytical).

### LC/ESI mass spectrometry

Liquid Chromatography Electrospray Ionization Mass Spectrometry (LC/ESI-MS) was performed on a 6210 LC-TOF spectrometer coupled to an HPLC system (Agilent Technologies). All solvents used were HPLC grade (Chromasolv, Sigma-Aldrich). Trifluoroacetic acid (TFA) was from Acros Organics (puriss., p.a.). Solvent A was 0.03% TFA in water, solvent B was 95% acetonitrile-5% water-0.03% TFA. Just before analysis NME1 samples (57 mM in sample buffer: 20 mM Tris-HCl pH 8.8, 5 mM MgCl_2_, 1 mM DTT) were diluted with sample buffer to a final concentration of 5.7 µM and 4 ml were injected for MS analysis. Protein samples were first desalted on a reverse phase-C8 cartridge (Zorbax 300SB-C8, 5 mm, 300 µm ID’5mm, Agilent Technologies) for 3 min at a flow rate of 50 ml/min with 100% solvent A and then eluted with 70% solvent B at a flow rate of 50 ml/min for MS detection. MS acquisition was carried out in positive ion mode in the 300- 3200 *m/z* range. MS spectra were acquired and the data processed with MassHunter workstation software (v. B.02.00, Agilent Technologies) and with GPMAW software (v. 7.00b2, Lighthouse Data, Denmark).

### Native mass spectrometry

Mass Spectrometry under native conditions was performed to detect NME1 species following incubation with either CoA, dephospho-CoA, AcCoA, SucCoA, or a mixture of both CoA and ATP (all from Sigma-Aldrich). NME1 (50 µM) was incubated with the ligands (5 to 2500 µM) in a buffer containing 250 mM ammonium acetate, 0.5 mM magnesium acetate and DTT 1 mM for 15 min at room temperature and subsequently loaded by a nanoflow platinum-coated borosilicate electrospary capillary (Thermo Electron SAS; Courtaboeuf, France) into a quadrupole time-of-flight mass spectrometer (nano-ESI-Q-TOF instrument, Q-TOF Ultima, Waters Corporation, Manchester, U.K.). The following instrumental parameters were used: capillary voltage = 1.2-1.3 kV, cone potential = 40 V, RF lens-1 potential = 40 V, RF lens-2 potential = 1 V, aperture-1 potential = 0 V, collision energy = 30-140 V, and microchannel plate (MCP) = 1900 V. The pressure in the collision cell was set to ∼2 × 10^−2^ mbar. All mass spectra were calibrated externally using a solution of cesium iodide (6 mg/mL in 50% isopropanol) and were processed with the Masslynx 4.0 software (Waters Corporation, Manchester, U.K.) and with Massign software package.

### In vitro CoA binding assay of the NME1 protein

CoA-pulldown was performed in the presence of purified WT NME1 or the T94D mutant. Three μg of WT or mutant GST-tagged or untagged NME1 were incubated overnight at 4°C (300 μl final volume) with 50 µl of pre-washed CoA-agarose resin (buffer: 20% glycerol, 3 mM MgCl_2_, 50 mM Hepes, 500 mM KCl, 1 mM DTT, 1 mini-tablet of EDTA-free protease inhibitor) in the presence or absence of ATP (Sigma-Aldrich). The enriched CoA-agarose was washed three times in the buffer and, after a final wash with PBS, was incubated for 1 h at 4°C in 50 μl of 5 mM free ligand (either CoA, NAD, FAD, ADP-ribose, ATP or dephospho-CoA; all from Sigma-Aldrich) or 50 μL H_2_O as a Control. Flow-through and CoA-agarose (eluted by free CoA) fractions were analyzed by western blotting to detect NME1 (by an anti-GST or anti-NME2 antibody) as well as histidine phosphorylation.

### Determination of nucleoside diphosphate kinase activity

NME kinase activity was determined spectrophotometrically with 4 mM ATP and 1 mM TDP as substrates using a coupled enzyme assay. ADP production was coupled by pyruvate kinase (160 U/mL) and lactate dehydrogenase (800 U/mL) to NADH oxidation using 4.5 mM Mg-acetate, 0.9 mM PEP and 0.45 mM NADH in 0.1 M triethanolamine buffer pH 7. The assay was started by addition of NME (20 ng wild-type or 7 µg mutant/1 mL) and changes in NADH redox state were followed at 340 nm and 25°C for 10 min. NME kinase activity was then inhibited with 1 mM acetyl-CoA and followed for another 10 min. Data were corrected for non-specific inhibitory effects of acetyl-CoA on the coupled enzyme system by running controls with creatine kinase (4 mM ATP and 20 mM creatine as substrates) with the same protocol.

### Fatty acid detection

Total fatty acids were extracted from the liver using a 1:2 (v/v) mixture of chloroform/methanol in the presence of 10 nmol of tridecanoic acid (Sigma-Aldrich #T0502) and 10 nmol of phosphatidylcholine (PC, C21:0/C21:0, Avanti #850370) as an internal standard. The liver samples were dissected into small pieces and ground in methanol prior to the addition of chloroform. After vigorous sonication and vortexing, the organic phase was extracted by biphasic separation generated by the addition of chloroform and 0.2% KCl in order to obtain a chloroform/methanol/aqueous ratio of 2:1:0.8 (v/v/v). The resulting bottom organic phase was dried and on-line derivatized to fatty acid methyl ester by a chloroform/methanol (1:2)-TMSH solution (Macherey-Nagel) and analyzed by gas chromatography-mass spectrometry (Agilent 5977A-7890B). The abundance of each fatty acid was calculated and normalized according to the internal standard and cell number or wet weight.

### High-fat diet treatment

The high-fat diet treatment was approved by the official ethics committee of the University Grenoble Alpes (ComEth, C2EA-12). WT and Nme2 ko mice (C57BL/6N- Nme2tm1.1(KOMP)Vlcg/MbpMmucd; MMRRC stock #: 048819-UCD) were fed with Normal diet (Safe A03, France) and High fat custom diet (SAFE® U8978 Version 19) for 6 weeks. The food and mice were weighed every 3-4 days.

### Detection of adenine nucleotides, CoA and acetyl-CoAs in liver

Adenine nucleotides, CoA and acetyl-CoAs were determined in protein-free extracts obtained by perchloric acid precipitation as follows. Freeze-clamped livers were homogenized in 0.5 N perchloric acid (2:1 v/w) in liquid nitrogen. After thawing at room temperature, 4 N perchloric acid (1:10 v/w) was added, and the homogenate was incubated on ice for 30 min and centrifuged (4000 g, 5 min, 4°C). The supernatant was neutralized by addition of 5 M K_2_CO_3_ and centrifuged again (4000 g, 5 min, 4°C). The metabolites were determined by HPLC (Varian 410, France) with RP-C18 column (Polaris C18-A 250 × 4.6, 5 μm, Varian, France; ref. A2000250R046) at a 1 mL/min flow rate at 30°C, as described in detail elsewhere^82, 83^. Briefly, to determine adenine nucleotides, the protein-free extract (75 mL premixed with 50 mL mobile phase) was separated in pyrophosphate buffer (28 mM, pH 5.75), and the detection was performed at 254 nm. The ATP, ADP, and AMP were eluted at 6.5, 8, and 14 min, respectively. To determine CoA and acetyl-CoA, the protein-free extract (75 mL premixed with 50 mL mobile phase) complemented with DTT (2 mM final) was separated in 100 mM monosodium phosphate, 75 mM sodium acetate, pH 4.6 combined with acetonitrile in the proportion 94:6 (v/v). The detection was performed at 259 nm. The CoA and acetyl-CoA were respectively eluted at 7, and 16 min. The elution peaks were integrated using the STAR software (Varian, France).

### Measurement of liver triglyceride content

Frozen liver fragments (50 mg) were digested in 0.15 mL of 3 M alcoholic potassium hydroxide (70°C, 2 hours) and the amount of liver triglycerides was measured using a Triglycerides kit (Erba Mannheim, Brno, Czech Republic, Ref: BLT00059, Lot 2107029), and spectroscopy to measure the samples’ absorbance.

### Histological analyses

The liver tissues were fixed in a formalin solution, neutral buffered at 10% (Sigma-Aldrich, Steinheim am Albuch, Germany) and paraffin-embedded. Four-micrometer sections of tissue were prepared. Hematoxylin-eosin (HE) staining was used for histopathological examination. The presence of steatosis was determined through a blind evaluation of the slides.

### HepG2 cell lines

Nme1 and Acly were knocked down in liver hepatocarcinoma cells (HepG2) using Lipofectamine RNAiMAX (Invitrogen #11668-019)-siRNA transfection (siCtr – Eurogentec SR-CL011-005). SiRNA targeting sequences for human Nme1 and Acly are reported in the key resources table.

### Transcriptome (RNAseq)

RNAseq overall design

Transcriptomic analysis was performed on mouse liver samples obtained from 4 conditions, corresponding to two genotypes, Nme2 WT (Nme2^+/+^) and KO (Nme2^-/-^), and two experimental diets, normal diet (ND) and high fat diet (HFD). For each condition, RNAseq was performed in 4 replicates corresponding to 4 individual mice.

### Protocols Extraction protocol

Total RNA was isolated and purified from the liver tissue by NucleoSpin RNA kit (Machery-Nagel- 740955.50). Four independent RNA extractions were performed for each condition.

### Library construction protocol

For each sample, one μg of RNA (RIN>8) was used for library preparation with the Illumina Stranded total RNA Prep Ligation with Ribo-Zero Plus kit (Illumina) according to the manufacturer’s instructions. Each library was quantified on Qubit with the Qubit® dsDNA HS Assay Kit (Life Technologies) and the size distribution was examined on the Fragment Analyzer with High Sensitivity NGS Fragment Analysis kit (Agilent).

### Library strategy

The libraries were then sequenced on the Illumina NextSeq 500 (paired-end 75) at the TGML Platform of Aix-Marseille University (France).

### Data Processing Pipeline Alignment

The sequenced reads from the raw sequence (.fastq files) were aligned on the UCSC mm10 genome using the STAR software (2.7.1a)^84^ to produce bam files.

### Counts

The aligned reads (.bam files) were counted using HTSeq framework (0.11.2)^85^ with options: -t exon -f bam -r pos –stranded=reverse -m intersection-strict –nonunique none

### Normalization and differential analysis

The normalization, pseudo-log transformation of the read counts, and the differential analyses were performed using the R software [R Core Team. R: A language and environment for statistical computing. Vienna, Austria: R Foundation for Statistical Computing, 2017], DESeq2 (1.22.2)^86, 87^ and SARTools^88^ packages.

### Grouping genes/features according to their quartile of expression

The features corresponding to non-coding transcripts and/or with zero counts were filtered out, in order to retain all non-zero count protein-coding genes from the NCBI Reference Sequence Database (RefSeq). The corresponding genes and features were ranked according to their DESeq2 normalized expression mean value in the samples corresponding to the 4 replicates of liver from wild-type mice under normal diet, and allocated into four subsets corresponding to the quartiles of this expression and exported as .bed files.

### Identification of up- and down-regulated genes and selection for heatmap representation

Supervised transcriptomic analyses were performed to identify genes significantly up- and down- regulated between two conditions using thresholds of a Student *t*-test *p*-value <0.01 and a fold change absolute value of 2.

The normalized, pseudo-log transformed and standardized read counts of the up- and down- regulated genes in wild type mice between those submitted to a high fat diet (HFD) and those with a normal diet (ND) were used to generate the heatmap presented in **Fig. 4a**.

### ChIPseq sequencing and analysis H3K9ac ChIP assay

ChIP assays for H3K9ac were carried out as previously described with minor modifications^69, 70^. Briefly, 50 µl of Dynabeads protein G (Thermo-fisher 10004D) were washed 3 times with blocking solution (5% BSA/PBS) and incubated in 500 µl overnight at 4°C with 5 µl of H3K9ac antibody. On the second day, liver tissue was homogenized in cold buffer (3 ml/each liver) containing 2.2 M sucrose, 10 mM Tris-HCl pH 7.5, 10 mM MgCl_2_, 10 mM sodium butyrate and protease inhibitor, filtered by hydrophilic gauze and centrifuged for 3 h at 100,000 g at 4°C. Nuclei in the pellet were washed in a buffer containing 0.5 M sucrose, 10 mM Tris-HCl pH 7.5, 10 mM MgCl_2_, 0.2% Triton X-100, 10 mM sodium butyrate, protease inhibitor, centrifuged for 1467 g at 4°C and resuspended in a buffer containing 1 M sucrose, 10 mM Tris-HCl, 10 mM MgCl_2_, 10 mM sodium butyrate (Sigma-Aldrich B-2503) and protease inhibitor. 45 µg of nuclei were digested with 0.75 µg of Nuclease S7 (Sigma-Aldrich, Cat#10107921001) for 15 min at 37 °C in 100 µl of a buffer containing 20 mM of Tris-HCl pH7.5, 5 mM CaCl_2_ to obtain mononucleosomes (146 bp). The reaction was stopped by adding 5 mM EDTA. Small aliquots of mononucleosome solutions were collected for input. Digested mononucleosomes were diluted with LSDB buffer (50 mM HEPES pH 7.0, 3 mM MgCl_2_, 500 mM KCl, 20% glycerol, protease cocktail inhibitor containing 10 mM sodium butyrate) to achieve the final KCl concentration of 350 mM and incubated with antibody-coupled beads overnight at 4°C for 16 hours. On the third day, the beads were washed four times with LSDB 350 mM KCl and once in TE buffer (10 mM Tris-HCl containing 1 mM EDTA, pH 8.0). ChIP samples were eluted in 150 µl of the buffer with SDS 1% for 20 min at 65°C. The DNA was purified from the eluted ChIP samples as well as from the input samples by phenol-chloroform extraction and ethanol precipitation.

### ChIPseq overall design

ChIPseq analysis was performed in mouse liver samples obtained from 4 conditions, corresponding to two genotypes, Nme2 WT (Nme2^+/+^) and KO (Nme2^-/-^), and two experimental diets, normal diet (ND) and high fat diet (HFD). For each condition, chromatin immunoprecipitation was performed twice in the respective livers of two independent mice using anti-H3K9ac antibody of mNase digested chromatin and both the immune-precipitated (chip) and input materials were sequenced.

### Protocols Extraction protocol

As previously reported, the DNA was purified from the eluted ChIP samples as well as from the input samples by phenol-chloroform extraction and ethanol precipitation.

### Library construction protocol

For sequencing, ChIP libraries were prepared using MicroPlex Library Preparation Kit v3 (Diagenode) according to manufacturer’s instructions. Each library was quantified on Qubit with Qubit® dsDNA HS Assay Kit (Life Technologies) and size distribution was examined on the Fragment Analyzer with High Sensitivity NGS Fragment Analysis kit (Agilent).

### Library strategy

The ChIP libraries were sequenced on a High-output flow cell (400M clusters) using the NextSeq® 500/550 High Output v2.5 150 cycles kit (Illumina), in paired-end 75/75nt mode, according to manufacturer’s instructions at the TGML Platform of Aix-Marseille University (France). Base calling was performed using RTA version 2.*

### Data Processing Pipeline

Data processing step: Trimming

The raw fastq files were processed by 5 prime trimming, keeping 30bp-length fragments, using fastx_trimmer [http://hannonlab.cshl.edu/fastx_toolkit/. Accessed 28 Feb. 2022.], with options - l 30 -Q33.

Data processing step: Alignment

The trimmed fastq files were aligned on the UCSC Mus_musculus mm10 genome using the Bowtie2 aligner^89^, with options –end-to-end, –no-mixed, –no-discordant.

Genome build: UCSC Mus_musculus mm10 genome

Processed data files format and content: big wig files (.bw) containing normalized integrated aligned read count signals.

### Selection of features corresponding to TSS and normalization

Raw ChIP counts corresponding to TSS +1500/-800bp of the RefSeq genes were computed using featureCounts^90^, with option -a mm10_pc_tss8001500.saf -F SAF -s 0 -Q 30 -T 8 -o mm10_pc_tss8001500_count_30.txt.

For each sample, scale factors were computed on these features using DESeq2^86, 87^ assuming that the global level of ChIP signal value (corresponding lysine acetylation in all genes promoters) should be equal in all samples.

The scale factors are given in the “Deseq2_sf “ table shown below.

| sample | sf | inv_sf | bw_file |
| --- | --- | --- | --- |
| normal_chip_nme2wt_m301 | 1 | 1,13 | normal_chip_nme2wt_m301_end-to-end_trim30_srt_mmq30_PESF_30_4_None_SF1.13.bw |
| normal_chip_nme2wt_m302 | 1 | 1,48 | normal_chip_nme2wt_m302_end-to-end_trim30_srt_mmq30_PESF_30_4_None_SF1.48.bw |
| hghfat_chip_nme2wt_m300 | 1 | 0,7 | hghfat_chip_nme2wt_m300_end-to-end_trim30_srt_mmq30_PESF_30_4_None_SF0.70.bw |
| hghfat_chip_nme2wt_m303 | 1 | 0,77 | hghfat_chip_nme2wt_m303_end-to-end_trim30_srt_mmq30_PESF_30_4_None_SF0.77.bw |
| normal_chip_nme2ko_m291 | 1 | 1,31 | normal_chip_nme2ko_m291_end-to-end_trim30_srt_mmq30_PESF_30_4_None_SF1.31.bw |
| normal_chip_nme2ko_m293 | 1 | 1,08 | normal_chip_nme2ko_m293_end-to-end_trim30_srt_mmq30_PESF_30_4_None_SF1.08.bw |
| hghfat_chip_nme2ko_m273 | 1 | 0,95 | hghfat_chip_nme2ko_m273_end-to-end_trim30_srt_mmq30_PESF_30_4_None_SF0.95.bw |
| hghfat_chip_nme2ko_m292 | 1 | 0,79 | hghfat_chip_nme2ko_m292_end-to-end_trim30_srt_mmq30_PESF_30_4_None_SF0.79.bw |

The bam signals were normalized and smoothed using bamCoverage (from deepTools suite^91^) with options: –binSize 4 –minMappingQuality 30 –normalizeUsing None –scaleFactor inv_sf where inv_sf is the inverse of the scaling factor previously computed

### ChIPseq Heatmaps

Repeat masker tagged loci GSAT_MM and SYNREP_MM as well as the Sfi1 locus (chr11:3126500- 3200500) and mitochondrial (chrM) genes were excluded from the analysis.

The normalized ChIP signals were converted into a 10bp bin matrix of the signal 1.5Kb upstream and downstream of protein-coding genes TSS, using computeMatrix (from deepTools suite^91^, with options reference-point –referencePoint TSS –binSize 10 –beforeRegionStartLength 1500 – afterRegionStartLength 1500 –sortRegions descend Heatmaps were generated using plotHeatmap (deepTools suite^91^ with options –colorMap YlOrRd –sortRegions descend).

### Highly expressed gene TSS profiles

The normalized ChIP signal were converted into a 10bp bin matrix of the signal 1.5Kb upstream and downstream 25% more expressed (see Mat & Meth RNA-seq) protein-coding genes TSS, using computeMatrix.

The ChIPseq profiles were generated using the computed computeMatrix outputs and a custom R script [R Core Team (2019). R: A language and environment for statistical computing.].

### Expression vs. Acetylation

The normalized ChIP signals were converted into a 25bp bin matrix of the signal 750bp upstream and 1500bp downstream protein-coding genes TSS, using computeMatrix.

For each gene and condition the mean signal over this region in all replicates was plotted.

**Fig. 4h**, the x-axis represents the log_2_ (fold changes) of the differential expression (DESeq2 normalized values) between HFD and ND respectively for Nme2 WT (upper panel) and KO (lower panel).

### Antibodies’ Dilutions for WB

Anti Nme2: 1/5000; Acc1 1/1000; P-Acc1 1/1000; AMPKα 1/1000, P-AMPKα 1/1000, Acly: 1/1000; ACSS2 1/1000; GAPDH 1/1000; β-Tubulin: 1/2000, α-Tubulin 1/2000, β-actin: 1/2000, Histidine 1P (SC1-1 clone) 1/1000, H3K9ac 1/1000, H3 1/5000.

## QUANTIFICATION AND STATISTICAL ANALYSIS

MS-based proteomics of the CoA bound proteins (**Fig. 1c-e** and **Fig. S1**): 4 testes from 2 adult male mice were the biological source of total testis CoA bound proteins, while 12 testes from 6 adult males were the source of fractionated spermatogenic cells CoA bound proteins.

NME1/2 WB in liver (**Fig. 3a**): (2 month mice); 2 *Nme2^+/+^* (2 females (F)), *2 Nme2^-/-^* (2 F).

CoA pull down liver-silver staining: (**Fig. 3b**): (2-3 month mice); 3 *Nme2^+/+^* (3 males (M)), 3 *Nme2^-/-^* (3M).

Fatty acid detection (**Fig. 3c,d**): the graphs show the average values and SEM. Liver from adult mice (2-3 months); 5 *Nme2^+/+^* (2 F, 3 M), 4 *Nme2^-/-^* (2 F, 2 M). Total Fatty Acids (**Fig. 3c**) **p value = 0.0021, different fatty acids species (**Fig. 3d**) ***p <0.001. Multiple unpaired t-test and correction for multiple comparison using the Hom-Sidak method.

Steatosis in liver (**Fig. 3e**): (2-4 month mice); 4 *Nme2^+/+^* in ND (normal diet) (2 F, 2 M) – steatosis was not detected in these liver samples; 5 *Nme2^-/-^* in ND (2 F, 3 M) - no steatosis; 6 *Nme2^+/+^* in HFD (high-fat diet) (1 F, 5 M) - no steatosis; 7 *Nme2^-/-^* in HFD mice (3 F, 4 M) – steatosis was detected in the 4 M, and initial steatosis in the 3 F.

Triglyceride detection in liver (**Fig. 3f**): (3-5 month mice); 4 *Nme2^+/+^* (2 F, 2 M) in ND, 4 *Nme2^-/-^* (2 F, 2 M) in ND, 10 *Nme2^+/+^* (3 F, 7 M) in HFD, 12 *Nme2^-/-^* (5 F, 7 M) in HFD. The graph shows the average values and SEM. ND: *Nme2^+/+^* vs *Nme2^-/-^,* *p value = 0.0286; HFD: *Nme2^+/+^* vs *Nme2^-/-^*,** p value = 0.0071. Mann-Whitney t-test.

RNA-seq (**Fig. 4a**): (2-4 months mice), 4 *Nme2^+/+^* in ND (4 M), 4 *Nme2^+/+^* in HFD (4 M), 4 *Nme2^-/-^* in ND (4 M), 4 *Nme2^-/-^* in HFD (4 M).

CoA and AcetylCoA detection in liver (**Fig. 4e**): (3-5 month mice), 9 *Nme2^+/+^* in ND (2 F, 7 M), 4 *Nme2^+/+^* in HFD (2 F, 2 M), 9 *Nme2^-/-^* in ND (2 F, 7 M), 5 *Nme2^-/-^* in HFD (2 F, 3 M). The graph shows the average values and SEM. CoA: *** p = 0.003 *Nme2^+/+^* (ND vs HFD), *** p = 0.007 *Nme2^-/-^* (ND vs HFD). AcCoA: *Nme2^-/-^* (ND vs HFD) ** P = 0.002 KO. 2-way Anova test (with diet and genotype as the two factors).

Chip-seq and H3K9ac WB (**Fig. 4f** and **d**): (3-5 month mice), 2 *Nme2^+/+^* in ND (2 M), 2 *Nme2^+/+^* in HFD (2 M), 2 *Nme2^-/-^* in ND (2 M); 4 *Nme2^-/-^* in HFD (2 M).

WB in HepG2: (**Fig. 5a**). The panel shows the experiment in triplicates. Relative intensity ACC1/β- actin, si*CTR* versus Si*Nme1* \*\**p*=0.0059; si*Nme1* versus si*Acly* \**p*=0.0180, one-way ANOVA test. WB in liver (**Fig. S13a**): (2-4 months mice); 3 *Nme2^+/+^* in ND (3 M), 3 *Nme2^-/-^* in ND (3 M), 4 *Nme2^+/+^* in HFD (4 M), 4 *Nme2^-/-^* in HFD (4 M).

ATP/ADP/AMP detection in liver (**Fig. S13b**): (3-5 month mice), 9 *Nme2^+/+^* in ND (2 F, 7 M), 4 *Nme2^+/+^* in HFD (2 F, 2 M), 9 *Nme2^-/-^* in ND (2 F, 7 M), 5 *Nme2^-/-^* in HFD (2 F, 3 M). The graph shows the average values and SEM.

## Notes

### Competing Interest Statement

The authors have declared no competing interest.

## REFERENCES

1. Wallace, M. & Metallo, C. M. Tracing insights into de novo lipogenesis in liver and adipose tissues. Semin Cell Dev Biol 108, 65–71 (2020).

2. Ameer, F., Scandiuzzi, L., Hasnain, S., Kalbacher, H. & Zaidi, N. De novo lipogenesis in health and disease. Metabolism 63, 895–902 (2014).

3. Lambert, J. E., Ramos-Roman, M. A., Browning, J. D. & Parks, E. J. Increased de novo lipogenesis is a distinct characteristic of individuals with nonalcoholic fatty liver disease. Gastroenterology 146, 726–735 (2014).

4. Smith, G. I. et al. Insulin resistance drives hepatic de novo lipogenesis in nonalcoholic fatty liver disease. J Clin Invest 130, 1453–1460 (2020).

5. Roumans, K. H. M. et al. Hepatic saturated fatty acid fraction is associated with de novo lipogenesis and hepatic insulin resistance. Nat Commun 11, 1891 (2020).

6. Kasper, P. et al. NAFLD and cardiovascular diseases: a clinical review. Clin Res Cardiol 110, 921–937 (2021).

7. Lee, Y. et al. Serial Biomarkers of De Novo Lipogenesis Fatty Acids and Incident Heart Failure in Older Adults: The Cardiovascular Health Study. J Am Heart Assoc 9, e014119 (2020).

8. Koundouros, N. & Poulogiannis, G. Reprogramming of fatty acid metabolism in cancer. Br J Cancer 122, 4–22 (2020).

9. Schug, Z. T., Vande Voorde, J. & Gottlieb, E. The metabolic fate of acetate in cancer. Nat Rev Cancer 16, 708–717 (2016).

10. Zaidi, N., Swinnen, J. V. & Smans, K. ATP-Citrate Lyase: A Key Player in Cancer Metabolism. Cancer Research 72, 3709–3714 (2012).

11. Ke, R., Xu, Q., Li, C., Luo, L. & Huang, D. Mechanisms of AMPK in the maintenance of ATP balance during energy metabolism: AMPK and ATP balance. Cell Biol Int 42, 384–392 (2018).

12. Boissan, M. & Lacombe, M.-L. Learning about the functions of NME/NM23: lessons from knockout mice to silencing strategies. Naunyn-Schmiedeberg’s Arch Pharmacol 384, 421–431 (2011).

13. Adam, K., Ning, J., Reina, J. & Hunter, T. NME/NM23/NDPK and Histidine Phosphorylation. IJMS 21, 5848 (2020).

14. Yu, B. Y. K. et al. Regulation of metastasis suppressor NME1 by a key metabolic cofactor coenzyme A. Redox Biol 44, 101978 (2021).

15. Zhang, S. et al. Long-chain fatty acyl coenzyme A inhibits NME1/2 and regulates cancer metastasis. Proc. Natl. Acad. Sci. U.S.A. 119, e2117013119 (2022).

16. Lv, L. & Lei, Q. Proteins moonlighting in tumor metabolism and epigenetics. Front. Med. 15, 383–403 (2021).

17. Goudarzi, A., Shiota, H., Rousseaux, S. & Khochbin, S. Genome-scale acetylation- dependent histone eviction during spermatogenesis. J Mol Biol 426, 3342–3349 (2014).

18. Goudarzi, A. et al. Dynamic Competing Histone H4 K5K8 Acetylation and Butyrylation Are Hallmarks of Highly Active Gene Promoters. Mol Cell 62, 169–180 (2016).

19. Shiota, H. et al. Nut Directs p300-Dependent, Genome-Wide H4 Hyperacetylation in Male Germ Cells. Cell Rep 24, 3477–3487.e6 (2018).

20. Pusch, W., Balvers, M., Weinbauer, G. F. & Ivell, R. The Rat Endozepine-Like Peptide Gene Is Highly Expressed in Late Haploid Stages of Male Germ Cell Development1. Biology of Reproduction 63, 763–768 (2000).

21. Boeri Erba, E., Signor, L. & Petosa, C. Exploring the structure and dynamics of macromolecular complexes by native mass spectrometry. Journal of Proteomics 222, 103799 (2020).

22. Cervoni, L. et al. Binding of Nucleotides to Nucleoside Diphosphate Kinase: A Calorimetric Study. Biochemistry 40, 4583–4589 (2001).

23. Chen, Y. et al. Adenosine Phosphonoacetic Acid is Slowly Metabolized by NDP Kinase. MC 1, 529–536 (2005).

24. Dumas, C. et al. X-ray structure of nucleoside diphosphate kinase. The EMBO Journal 11, 3203–3208 (1992).

25. Schneider, B., Xu, Y. W., Janin, J., Véron, M. & Deville-Bonne, D. 3′-Phosphorylated Nucleotides Are Tight Binding Inhibitors of Nucleoside Diphosphate Kinase Activity. Journal of Biological Chemistry 273, 28773–28778 (1998).

26. Giraud, M.-F., Georgescauld, F., Lascu, I. & Dautant, A. Crystal Structures of S120G Mutant and Wild Type of Human Nucleoside Diphosphate Kinase A in Complex with ADP. J Bioenerg Biomembr 38, 261–264 (2006).

27. Janin, J. et al. Three-dimensional structure of nucleoside diphosphate kinase. J Bioenerg Biomembr 32, 215–225 (2000).

28. Bourdais, J. et al. Cellular Phosphorylation of Anti-HIV Nucleosides. Journal of Biological Chemistry 271, 7887–7890 (1996).

29. Schneider, B. et al. Pre-steady State of Reaction of Nucleoside Diphosphate Kinase with Anti-HIV Nucleotides. Journal of Biological Chemistry 273, 11491–11497 (1998).

30. Morera, S., Chiadmi, M., LeBras, G., Lascu, I. & Janin, J. Mechanism of phosphate transfer by nucleoside diphosphate kinase: X-ray structures of the phosphohistidine intermediate of the enzymes from Drosophila and Dictyostelium. Biochemistry 34, 11062–11070 (1995).

31. Postel, E. H., Zou, X., Notterman, D. A. & La Perle, K. M. D. Double knockout Nme1/Nme2 mouse model suggests a critical role for NDP kinases in erythroid development. Mol Cell Biochem 329, 45–50 (2009).

32. Carrer, A. et al. Impact of a High-fat Diet on Tissue Acyl-CoA and Histone Acetylation Levels. J Biol Chem 292, 3312–3322 (2017).

33. Duarte, J. A. G. et al. A high-fat diet suppresses de novo lipogenesis and desaturation but not elongation and triglyceride synthesis in mice. J Lipid Res 55, 2541–2553 (2014).

34. Lundsgaard, A.-M. et al. Mechanisms Preserving Insulin Action during High Dietary Fat Intake. Cell Metabolism 29, 50–63.e4 (2019).

35. Newman, J. C. et al. Ketogenic Diet Reduces Midlife Mortality and Improves Memory in Aging Mice. Cell Metab 26, 547–557.e8 (2017).

36. Ozaki, M. Cellular and molecular mechanisms of liver regeneration: Proliferation, growth, death and protection of hepatocytes. Seminars in Cell & Developmental Biology 100, 62–73 (2020).

37. Hsieh, W.-C. et al. Glucose starvation induces a switch in the histone acetylome for activation of gluconeogenic and fat metabolism genes. Molecular Cell 82, 60–74.e5 (2022).

38. Mendoza, M. et al. Enzymatic transfer of acetate on histones from lysine reservoir sites to lysine activating sites. Sci. Adv. 8, eabj5688 (2022).

39. Gates, L. A. et al. Acetylation on histone H3 lysine 9 mediates a switch from transcription initiation to elongation. Journal of Biological Chemistry 292, 14456–14472 (2017).

40. Martin, B. J. E. et al. Transcription shapes genome-wide histone acetylation patterns. Nat Commun 12, 210 (2021).

41. Dankel, S. N. et al. Hepatic Energy Metabolism Underlying Differential Lipidomic Responses to High-Carbohydrate and High-Fat Diets in Male Wistar Rats. The Journal of Nutrition 151, 2610–2621 (2021).

42. Gao, X. et al. Acetate functions as an epigenetic metabolite to promote lipid synthesis under hypoxia. Nat Commun 7, 11960 (2016).

43. Zhao, S. et al. Dietary fructose feeds hepatic lipogenesis via microbiota-derived acetate. Nature 579, 586–591 (2020).

44. Zhang, H., Yang, Z., Shen, Y. & Tong, L. Crystal Structure of the Carboxyltransferase Domain of Acetyl-Coenzyme A Carboxylase. Science 299, 2064–2067 (2003).

45. Chow, J. D. Y. et al. Genetic inhibition of hepatic acetyl-CoA carboxylase activity increases liver fat and alters global protein acetylation. Mol Metab 3, 419–431 (2014).

46. Galdieri, L. & Vancura, A. Acetyl-CoA carboxylase regulates global histone acetylation. J Biol Chem 287, 23865–23876 (2012).

47. Rios Garcia, M., et al. Acetyl-CoA Carboxylase 1-Dependent Protein Acetylation Controls Breast Cancer Metastasis and Recurrence. Cell Metab 26, 842–855.e5 (2017).

48. Horie, S., Isobe, M. & Suga, T. Changes in CoA Pools in Hepatic Peroxisomes of the Rat, under Various Conditions. The Journal of Biochemistry 99, 1345–1352 (1986).

49. 49. Williamson, J. R. & Corkey, B. E. [23] Assay of citric acid cycle intermediates and related compounds—Update with tissue metabolite levels and Intracellular Distribution. in Methods in Enzymology vol. 55 200–222 (Elsevier, 1979).

50. Leonardi, R., Zhang, Y., Rock, C. & Jackowski, S. Coenzyme A: Back in action. Progress in Lipid Research 44, 125–153 (2005).

51. Idell-Wenger, J. A., Grotyohann, L. W. & Neely, J. R. Coenzyme A and carnitine distribution in normal and ischemic hearts. J Biol Chem 253, 4310–4318 (1978).

52. Taniguchi, Y. et al. Quantifying E. coli proteome and transcriptome with single-molecule sensitivity in single cells. Science 329, 533–538 (2010).

53. Beck, M. et al. The quantitative proteome of a human cell line. Mol Syst Biol 7, 549 (2011).

54. Schwanhäusser, B. et al. Global quantification of mammalian gene expression control. Nature 473, 337–342 (2011).

55. Mitsui, Y. & Schneider, E. L. Relationship between cell replication and volume in senescent human diploid fibroblasts. Mechanisms of Ageing and Development 5, 45–56 (1976).

56. Postel, E. H. Multiple biochemical activities of NM23/NDP kinase in gene regulation. J Bioenerg Biomembr 35, 31–40 (2003).

57. Puts, G. S., Leonard, M. K., Pamidimukkala, N. V., Snyder, D. E. & Kaetzel, D. M. Nuclear functions of NME proteins. Lab Invest 98, 211–218 (2018).

58. Postel, E. H., Weiss, V. H., Beneken, J. & Kirtane, A. Mutational analysis of NM23- H2/NDP kinase identifies the structural domains critical to recognition of a c-myc regulatory element. Proc. Natl. Acad. Sci. U.S.A. 93, 6892–6897 (1996).

59. Langer, M. R., Fry, C. J., Peterson, C. L. & Denu, J. M. Modulating Acetyl-CoA Binding in the GCN5 Family of Histone Acetyltransferases. Journal of Biological Chemistry 277, 27337– 27344 (2002).

60. Ngo, L., Brown, T. & Zheng, Y. G. Bisubstrate inhibitors to target histone acetyltransferase 1. Chem Biol Drug Des 93, 865–873 (2019).

61. Poux, A. N., Cebrat, M., Kim, C. M., Cole, P. A. & Marmorstein, R. Structure of the GCN5 histone acetyltransferase bound to a bisubstrate inhibitor. Proc. Natl. Acad. Sci. U.S.A. 99, 14065– 14070 (2002).

62. Tanner, K. G., Langer, M. R. & Denu, J. M. Kinetic Mechanism of Human Histone Acetyltransferase P/CAF. Biochemistry 39, 11961–11969 (2000).

63. Wapenaar, H. et al. Enzyme kinetics and inhibition of histone acetyltransferase KAT8. European Journal of Medicinal Chemistry 105, 289–296 (2015).

64. Albe, K. R., Butler, M. H. & Wright, B. E. Cellular concentrations of enzymes and their substrates. Journal of Theoretical Biology 143, 163–195 (1990).

65. Snider, N. T. et al. Energy determinants GAPDH and NDPK act as genetic modifiers for hepatocyte inclusion formation. Journal of Cell Biology 195, 217–229 (2011).

66. Liu, X. et al. Acetate Production from Glucose and Coupling to Mitochondrial Metabolism in Mammals. Cell 175, 502–513.e13 (2018).

67. Xu, Y.-W., Moréra, S., Janin, J. & Cherfils, J. AlF _3_ mimics the transition state of protein phosphorylation in the crystal structure of nucleoside diphosphate kinase and MgADP. Proc. Natl. Acad. Sci. U.S.A. 94, 3579–3583 (1997).

68. Perez-Riverol, Y. et al. The PRIDE database and related tools and resources in 2019: improving support for quantification data. Nucleic Acids Res 47, D442–D450 (2019).

69. Barral, S. et al. Histone Variant H2A.L.2 Guides Transition Protein-Dependent Protamine Assembly in Male Germ Cells. Molecular Cell 66, 89–101.e8 (2017).

70. Buchou, T., et al. Purification and Analysis of Male Germ Cells from Adult Mouse Testis. in HDAC/HAT Function Assessment and Inhibitor Development (ed. Krämer, O. H.) vol. 1510 159–168 (Springer New York, 2017).

71. Casabona, M. G., Vandenbrouck, Y., Attree, I. & Couté, Y. Proteomic characterization of Pseudomonas aeruginosa PAO1 inner membrane. Proteomics 13, 2419–2423 (2013).

72. Bouyssié, D. et al. Proline: an efficient and user-friendly software suite for large-scale proteomics. Bioinformatics 36, 3148–3155 (2020).

73. Couté, Y., Bruley, C. & Burger, T. Beyond Target–Decoy Competition: Stable Validation of Peptide and Protein Identifications in Mass Spectrometry-Based Discovery Proteomics. Anal. Chem. 92, 14898–14906 (2020).

74. Wieczorek, S., et al. DAPAR & ProStaR: software to perform statistical analyses in quantitative discovery proteomics. Bioinformatics 33, 135–136 (2017).

75. Kabsch, W. XDS. Acta Crystallogr D Biol Crystallogr 66, 125–132 (2010).

76. Vonrhein, C. et al. Data processing and analysis with the *autoPROC* toolbox. Acta Crystallogr D Biol Crystallogr 67, 293–302 (2011).

77. Evans, P. R. & Murshudov, G. N. How good are my data and what is the resolution? Acta Crystallogr D Biol Crystallogr 69, 1204–1214 (2013).

78. McCoy, A. J., et al. *Phaser* crystallographic software. J Appl Crystallogr 40, 658–674 (2007).

79. Mortenson, D. E. et al. “Inverse Drug Discovery” Strategy To Identify Proteins That Are Targeted by Latent Electrophiles As Exemplified by Aryl Fluorosulfates. J. Am. Chem. Soc. 140, 200–210 (2018).

80. Emsley, P., Lohkamp, B., Scott, W. G. & Cowtan, K. Features and development of *Coot*. Acta Crystallogr D Biol Crystallogr 66, 486–501 (2010).

81. Fuhs, S. R. et al. Monoclonal 1- and 3-Phosphohistidine Antibodies: New Tools to Study Histidine Phosphorylation. Cell 162, 198–210 (2015).

82. Gratia, S. et al. Inhibition of AMPK signalling by doxorubicin: at the crossroads of the cardiac responses to energetic, oxidative, and genotoxic stress. Cardiovascular Research 95, 290– 299 (2012).

83. Shurubor, Y. et al. Determination of Coenzyme A and Acetyl-Coenzyme A in Biological Samples Using HPLC with UV Detection. Molecules 22, 1388 (2017).

84. Dobin, A. et al. STAR: ultrafast universal RNA-seq aligner. Bioinformatics 29, 15–21 (2013).

85. Anders, S., Pyl, P. T. & Huber, W. HTSeq--a Python framework to work with high- throughput sequencing data. Bioinformatics 31, 166–169 (2015).

86. Anders, S. & Huber, W. Differential expression analysis for sequence count data. Genome Biol 11, R106 (2010).

87. Love, M. I., Huber, W. & Anders, S. Moderated estimation of fold change and dispersion for RNA-seq data with DESeq2. Genome Biol 15, 550 (2014).

88. Varet, H., Brillet-Guéguen, L., Coppée, J.-Y. & Dillies, M.-A. SARTools: A DESeq2- and EdgeR-Based R Pipeline for Comprehensive Differential Analysis of RNA-Seq Data. PLoS ONE 11, e0157022 (2016).

89. Langmead, B. & Salzberg, S. L. Fast gapped-read alignment with Bowtie 2. Nat Methods 9, 357–359 (2012).

90. Liao, Y., Smyth, G. K. & Shi, W. featureCounts: an efficient general purpose program for assigning sequence reads to genomic features. Bioinformatics 30, 923–930 (2014).

91. Ramírez, F., Dündar, F., Diehl, S., Grüning, B. A. & Manke, T. deepTools: a flexible platform for exploring deep-sequencing data. Nucleic Acids Research 42, W187–W191 (2014).

